# CochleaNet: deep learning-based image analysis for cochlear connectomics and gene therapy

**DOI:** 10.1101/2025.11.16.688700

**Authors:** Lennart Roos, Aleyna Miraç Diniz, Elisabeth Koert, Martin Schilling, Mara Uhl, Anupriya Thirumalai, Mostafa Aakhte, Kathrin Kusch, Jan Huisken, Tobias Moser, Constantin Pape

**Author notes:** equal contribution (Lennart Roos, Aleyna Miraç Diniz).

## Abstract

With the emergence of gene and optogenetic therapies targeting deafness, the comprehensive analysis of the molecular anatomy and physiology of the cochlea has become ever more important. Here, we introduce CochleaNet, a deep learning-based framework to analyze volumetric imaging data obtained by light-sheet microscopy of decalcified, cleared and fluorescently labeled cochleae. CochleaNet covers the workflow from reconstruction of the cochlea to segmentation of inner hair cells, spiral ganglion neurons and their afferent synapses, to analyzing the expression of gene therapy products. We validated CochleaNet by comparison to manual image analysis. Trained on high isotropic resolution mouse data, CochleaNet was also applicable to the cochlea of the gerbil, another relevant animal model, and lower-resolution mouse data from a commercially available microscope. We conclude that the combination of light-sheet microscopy and image analysis with CochleaNet paves the way for rapid and reliable quantification of cochlear molecular anatomy and preclinical gene therapy outcomes.

## Introduction

Multiscale photonic imaging of complete organs with subcellular resolution enables a new understanding of their morphology, physiology and pathology. Tissue clearing and light-sheet fluorescence microscopy (LSFM) are instrumental in these efforts (Weiss *et al*, 2021; Keller & Dodt, 2012; Ertürk *et al*, 2012; Renier *et al*, 2014; Santi, 2011). Recent advances have enabled the acquisition of large datasets, with applications ranging from developmental cellular atlases of small organisms (Lange *et al*, 2024) to studies of brain morphology and connectomics (Kaltenecker *et al*, 2024; Cai *et al*, 2019; Glaser *et al*, 2025). Sensory organs such as the ear and eye have attracted substantial interest for multiscale photonic imaging in fundamental, translational and clinical research (Santi, 2011; Ringel *et al*, 2021). Research on cochlear anatomy and physiology has heavily employed X-ray imaging <u>(Shibata und Nagano 1996; Rau et al. 2006; Bartels et al. 2013)</u>, LSFM (e.g. Santi *et al*, 2022; Keppeler *et al*, 2021; Moatti *et al*, 2020; Hutson *et al*, 2021; Cai *et al*, 2025; Aakhte *et al*, 2025), confocal and super-resolution microscopy (*e.g.* Kurima *et al*, 2015; Meyer *et al*, 2009; Neef *et al*, 2018; Kapoor *et al*, 2025), thereby covering the relevant length scales in separate experiments with complementary photonic techniques. Innovations in imaging techniques have increased resolution and extended their scope of investigation. For example, super-resolution microscopy of the cochlea has moved from resolving microclusters of proteins (e.g. Neef *et al*, 2018) to studying single proteins (Kapoor *et al*, 2025), complementing electron microscopy (EM) (e.g. Matsubara *et al*, 1996; Michanski *et al*, 2023). Moreover, the field has exploited the multiscale capability of X-ray and LSFM for imaging the cochlea from the whole organ down to the subcellular level (Schaeper *et al*, 2025; Aakhte *et al*, 2025).

In parallel, EM, the traditional method for imaging subcellular structures, has been extended to larger volumes including single hair cells or even multicellular stretches of the organ of Corti (e.g. Hua *et al*, 2021; Michanski *et al*, 2019; Payne *et al*, 2021), advancing our understanding of cochlear connectomics and hair cell ultrastructure, including their afferent synapses and their different functional states. The resulting volumetric datasets are huge, and their analysis remains a daunting task. This has motivated the development of machine learning-based image analysis, e.g. for mitochondria in serial block face EM (Liu *et al*, 2022; Karagulyan *et al*, 2025; Buswinka *et al*, 2023a) and ribbon synapses in electron tomography (Muth *et al*, 2025).

Like for volumetric EM, high-resolution photonic imaging of organs generates large volumetric datasets, posing a challenge to image processing and analysis (Schaeper *et al*, 2025; Aakhte *et al*, 2025). First, organs are imaged in tiles, which must be stitched to assemble the complete volume. For data handling and stitching, multiple standardized and scalable software tools are available (*e.g.*, Hörl *et al*, 2019). Novel methods scale to petabyte-sized data (Ruan *et al*, 2024). Further processing often includes denoising (Weigert *et al*, 2018) or other artifact removal steps (Peng *et al*, 2025). Next, addressing specific research questions requires a dedicated automated data analysis workflow. While the exact analysis steps may vary, they typically rely on segmentation to identify and localize cells or other structures of interest. Segmentation enables subsequent quantification, such as counting, morphological characterization, or spatial distribution analysis. Modern segmentation approaches are predominantly based on deep learning and the U-Net architecture (Falk *et al*, 2019), which has demonstrated superior performance for this task. The image analysis field has made significant progress in creating pretrained models that address cell segmentation tasks in diverse imaging modalities (Stringer *et al*, 2021; Schmidt *et al*, 2018; Archit *et al*, 2025). However, these models struggle with specific staining to target cellular and subcelluar structures, as well as efficient processing of large volumes such a LSFM data of the cochlea <u>(Keppeler et al. 2021)</u>. Consequently, more targeted approaches have been proposed for LSFM (Lange *et al*, 2024; Buswinka *et al*, 2023b; Malin-Mayor *et al*, 2023; Cai *et al*, 2025).

Yet, a solution for analyzing different cochlear cell types and subcellular structures, such as synapses, in LSFM data that is scalable and extensible to other cell types remains missing. Such a solution is needed for fundamental and translational research on the cochlea and other whole organs imaged in LSFM. For example, it would be instrumental for deciphering cochlear connectomics that involves i) afferent innervation of different types (I and II) and subtypes (I_a,b,c_) of spiral ganglion neurons (SGNs) that form heterogeneous afferent synapses with hair cells as well as ii) efferent innervation of SGNs and hair cells by brainstem neurons (Shrestha & Goodrich, 2019; Moser *et al*, 2023; Fuchs & Lauer, 2019). Moreover, major efforts are underway to preclinically develop gene therapeutic and optogenetic approaches for hearing restoration (Kleinlogel *et al*, 2020; Wolf *et al*, 2022; Petit *et al*, 2023; Carlson *et al*, 2025) and have paved the way for clinical proof of concept for gene therapy of otoferlin-related synaptic deafness (Lv *et al*, 2024; Qi *et al*, 2024, 2025). Advanced LSFM and deep learning-based image analysis would greatly enhance the efficiency of analyzing the expression and subcellular distribution of transgenic proteins in the preclinical development of such therapies.

Here, we present CochleaNet, a tool for the comprehensive analysis of cochleae imaged in LSFM. CochleaNet implements standardized image preprocessing, segmentation of SGNs and inner hair cells (IHCs), and detection of afferent ribbon synapses formed by IHCs and SGNs, based on dedicated deep learning models. The models were trained on newly annotated data, and we implemented efficient procedures to apply them to whole intact cochleae imaged by advanced LSFM (Aakhte *et al*, 2025). The isotropic sub-micrometer resolution of this custom LSFM facilitated robust cell segmentation and synapse detection in the complex cochlear morphology. We integrated the models into a unified data processing and analysis pipeline (Figure 1). Employing CochleaNet, we provided a comprehensive account of the number and tonotopic organization of SGNs, IHCs, and afferent ribbon synapses in the intact mouse cochlea. Moreover, we differentiated SGN subtypes across the cochlea based on additional fluorescent staining. We demonstrated the utility of CochleaNet for preclinical evaluation of gene and optogenetic therapies, highlighting how standardized LSFM and image analysis can facilitate the optimization of viral vectors, promoters and transgenes for gene therapies. Moreover, we demonstrated the utility of CochleaNet for the larger cochleae of gerbils and marmosets as well as for lower resolution anisotropic LSFM data. CochleaNet is available at https://github.com/computational-cell-analytics/cochlea-net.

**Figure 1.**
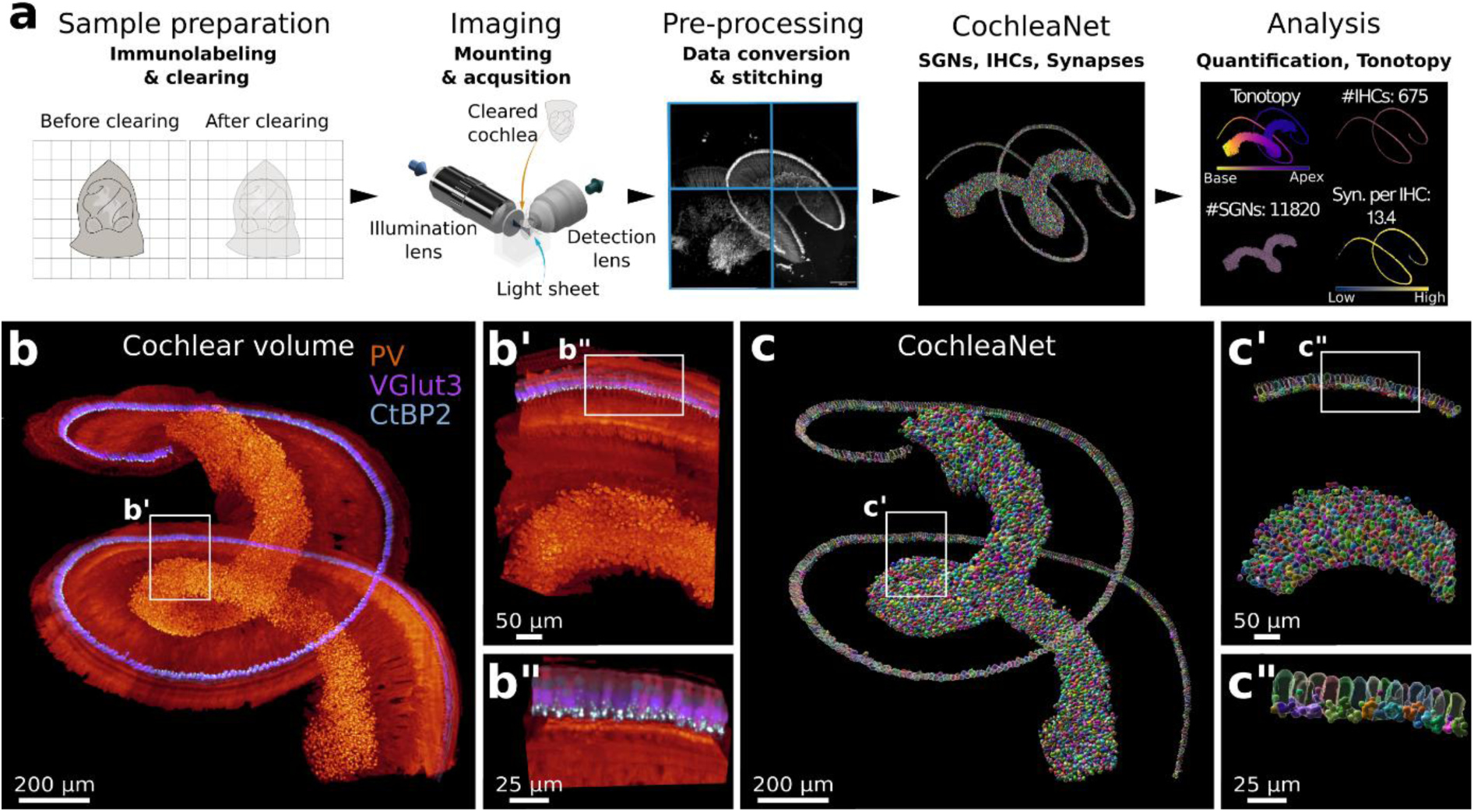
CochleaNet enables multiscale analysis of cochlear morphology. **a**, Schematic overview of the workflow from sample preparation, to imaging, to data pre-processing, to prediction with CochleaNet, and to subsequent data analysis. **b-b’’**, Intact mouse cochlea immunolabeled for parvalbumin (PV, red: labels spiral ganglion neuron (SGN) somata and neurites as well as inner hair cells (IHCs)), vesicular glutamate transporter 3 (Vglut3, blue: labels IHCs), and C-terminal Binding Protein 2 (CtBP2, white, staining RIBEYE of the presynaptic IHC ribbons). **c-c’’**, CochleaNet segmentation of SGNs based on PV, IHCs based on Vglut3 and ribbon synapse detections based on CtBP2, from three dedicated deep neural networks.

## Results

### Accurate quantification of neurosensory cell types and synapses in the mouse cochlea

We trained and evaluated three deep neural networks, collectively referred to as CochleaeNet, for specific analysis tasks. To this end, we first labelled, cleared and imaged four whole mouse cochleae using an advanced light-sheet microscope with isotropic sub-micron resolution (Figure 1, Methods) (Aakhte *et al*, 2025). We trained the network for SGN segmentation based on a parvalbumin (PV) staining and annotations of SGN somata forming the spiral ganglion in the Rosenthal’s canal (Eybalin & Ripoll, 1990). Note that PV staining also labels SGN neurites and IHCs (Figure 1). Here, we addressed SGN soma and IHC segmentation with two separate networks (Figure 1). We identify segmentation of peripheral SGN neurites, connecting SGN somata to IHCs, as a future task. Training the network for IHC segmentation was based on the IHC-specific labeling for vesicular glutamate transporter 3 (Vglut3) (Seal *et al*, 2008; Ruel *et al*, 2008). Finally, the detection of ribbon synapses was based on staining for C-terminal Binding Protein 2 (CtBP2) which labels ribbons and nuclei (Zenisek *et al*, 2004; Khimich *et al*, 2005). CtBP2 is a transcriptional co-repressor localized to nuclei and is homologous to the B-domain RIBEYE (Schmitz *et al*, 2000), which is the essential component of the ribbon (Maxeiner *et al*, 2016; Jean *et al*, 2018).

The training data consisted of 56 cochlear sub-volumes which were annotated by three experts semi-automatically. For SGNs (in PV) and IHCs (in Vglut3) this involved initial segmentation results from CellPose (Stringer *et al*, 2021) and µSAM (Archit *et al*, 2025) that were corrected manually in napari. For ribbon synapses, intensity-based detection of CtBP2 immunofluorescence spots was performed in the Imaris software (Meyer *et al*, 2009), followed by manual removal of false positive detections.

The three deep neural networks were implemented according to the 3D U-Net architecture (Falk *et al*, 2019). The segmentation networks for SGNs and IHCs were trained to predict foreground probabilities and distances to cell centroids and boundaries. These predictions were then post-processed to obtain a segmentation of individual cells using a watershed-based procedure (Archit *et al*, 2025). To exclude cells from adjacent cochlear tissues or other structures, a local density filter was applied to the SGN and IHC segmentations (Methods). The synapse detection network predicted a heightmap centered on synapses, extending a recent spot detection method (Jeremias & Pape, 2025) to volumetric data. Synapses were then identified with a local maxima filter. The detections were mapped to the closest segmented IHC within a maximum of three micrometers of its cell border. Using this methodology, we processed the image data of four entire cochleae (Ext. Data Table 1 and 2) for further validation and analysis. The segmentation results are visualized in Figure 1 and Ext. Data Figure 1.

To comprehensively compare and validate segmentation results, three experts manually annotated the center positions of all cells across twelve image planes of the four cochleae, for SGNs and IHCs respectively. We then created consensus annotations by matching these independent expert annotations against each other and retaining those that closely matched. We compared CochleaNet to state-of-the-art cell segmentation methods using these annotations: µSAM (Archit *et al*, 2025), CellPose (Stringer *et al*, 2021), Spiner (Cai *et al*, 2025), and StarDist (Schmidt *et al*, 2018) for SGNs as well as µSAM and CellPose for IHCs. These pretrained models were not specifically trained on our data, except for StarDist. The comparison was run on cochlear sub-volumes covering the annotated image planes and the segmented objects were matched to manual cell annotations for evaluation. We found that none of the other methods achieved IHC or SGN segmentation results of sufficient quality (Figure 2a and b, Ext. Data Figure 2). We assume that the specific intensity profiles and cell morphologies of IHCs and SGNs deviate from the training corpus of these established models, demonstrating the need for specialized solutions such as CochleaNet for analyzing complex volumetric imaging data.

**Figure 2:**
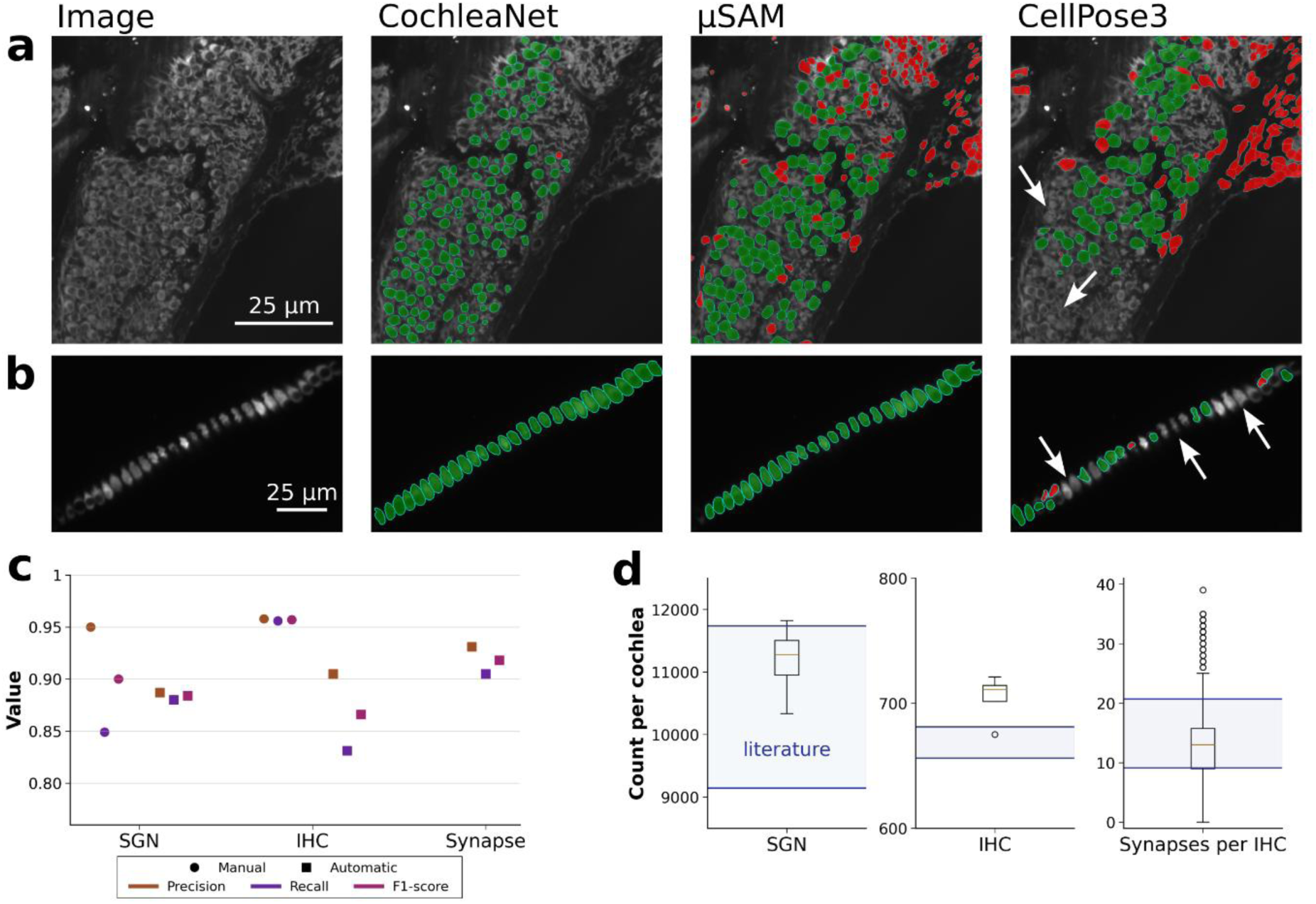
CochleaNet reliably segments SGNs, IHCs, and ribbon synapses in the mouse cochlea. **a**, SGN segmentation results of CochleaNet, µSAM, and CellPose3. White outlines indicate individual segmented cells, green overlays correctly segmented cells (true positives), red overlays wrongly segmented cells (false positives), white arrows indicate exemplary locations of missed cells (false negatives). **b**, IHC segmentation results of CochleaNet, µSAM, and CellPose3. **c**, Quantitative evaluation of SGN, IHC, and ribbon synapse predictions from CochleaNet based on comparison with expert annotations. For SGNs and IHCs, the consensus across annotations from three experts was used, and comparisons of individual expert annotations with the consensus were used to estimate the manual variability of the respective annotation tasks. We did not perform three independent annotations for the synapses due to the semi-automatic nature of synapse annotation. **d**, Number of SGNs and IHCs per cochlea as well as ribbon synapses per IHC derived from four cochleae, including reference values from literature. Circles indicate outliers not contained within the whiskers of the box plot. For synapses, the outliers likely result from multiple IHCs wrongly merged into a single object.

To validate the synapse detection, we annotated seven additional sub-volumes from two cochleae in the same manner as the respective synapse training data. We then applied CochleaNet to the four whole cochleae and evaluated the predictions for SGNs, IHCs, and synapses against the respective annotations, using the annotations for SGNs and IHCs described in the previous paragraph. We computed precision, recall, and F1-score based on true positives, false positives, and false negatives derived from these matches. We found that the predictions were accurate, with an F1-score of 0.884 for SGNs, 0.866 for IHCs, and 0.918 for synapses. For SGNs and IHCs, we also evaluated the independent expert annotations against the consensus to estimate annotator variability. The F1-score obtained for annotator variability was 0.9 for SGNs and 0.957 for IHCs (Figure 2c, Methods). Hence, the error rate of CochleaNet exceeded the variability in manual annotations for SGNs only marginally, likely because 3D annotation of SGNs is also challenging for humans. Annotation variability for IHCs was lower than CochleaNet’s error rate, likely because there are fewer IHCs, which can be more effectively inspected manually.

We then employed CochleaNet to quantify the morphology of the four cochleae in terms of their numbers of SGNs, IHCs, and synapses. On average, they contained 11,176 ± 542 SGNs, 705 ± 18 IHCs, and 11.8 ± 6.9 ribbon synapses per IHC, largely matching published data (Figure 2d, Ext. Data Table 3). These estimates tended to be at the upper edge of published data or larger. We argue that this points to the LSFM analysis being more comprehensive and less prone to error than in most previous studies as they i) cover all three key components of afferent cochlear connectivity and ii) do not miss cells or synapses due to microdissection of tissue or iii) rely on extrapolation as required when subsampling organ of Corti or spiral ganglion by stacks of confocal sections.

### Tonotopic mapping of IHCs, SGNs and afferent synapses in the mouse cochlea

The mammalian cochlea shows a frequency (tonotopic) map of the basilar membrane as a result of sound frequency decomposition by active and passive micromechanics (review in: Robles & Ruggero, 2001; Fettiplace, 2017). High frequency sounds best vibrate the stiffer and narrower basal end of the basilar membrane, while the softer and wider apex is best stimulated by low frequency sounds, which is typically described by the “Greenwood function” of the specific cochlea (review in: Greenwood, 1996). Interestingly, the innervation density of IHCs has been shown to be highest in the most sound sensitive midcochlear range probably reflecting optimal neural sampling of behaviorally most relevant signals (review in: Meyer & Moser, 2010; Rutherford & Moser, 2016).

Here, we analyzed the tonotopic distribution of SGNs (somata), IHCs, and ribbon synapses based on the CochleaNet segmentations. We first determined the run length from the base to the apex for the row of IHCs and for the spiral ganglion (Methods). We then fit Greenwood functions to the run length of IHCs and SGNs (Figure 3a, Methods). Next, we plotted the number of synapses per IHC as running average for the octaves along the tonotopic axis (Figure 3c). We found a rather invariant complement of ∼11-15 synapses per IHCs within the tonotopic range of 2-64 kHz, with lower values for low (< 2kHz) and high (>64 kHz) frequencies. For a comparison of our analysis to previous estimates of the tonotopic distribution of synapse number per IHC, we plotted data obtained by confocal imaging of microdissected fragments of the organ (Meyer *et al*, 2009) compared to our results (Ext. Data Figure 3a). We note that the total number of synapses per tonotopic range is even greater in the mid-frequency range of the cochlea because of the prevalence of staggered IHCs. Previously described by confocal microscopy (Yin *et al*, 2014) and EM (Liu *et al*, 2022), staggered IHCs were segmented by CochleaNet in most cases. We suspect that some of the outliers in Figure 2d result from rare failures of CochleaNet to faithfully segment staggered IHCs.

**Figure 3:**
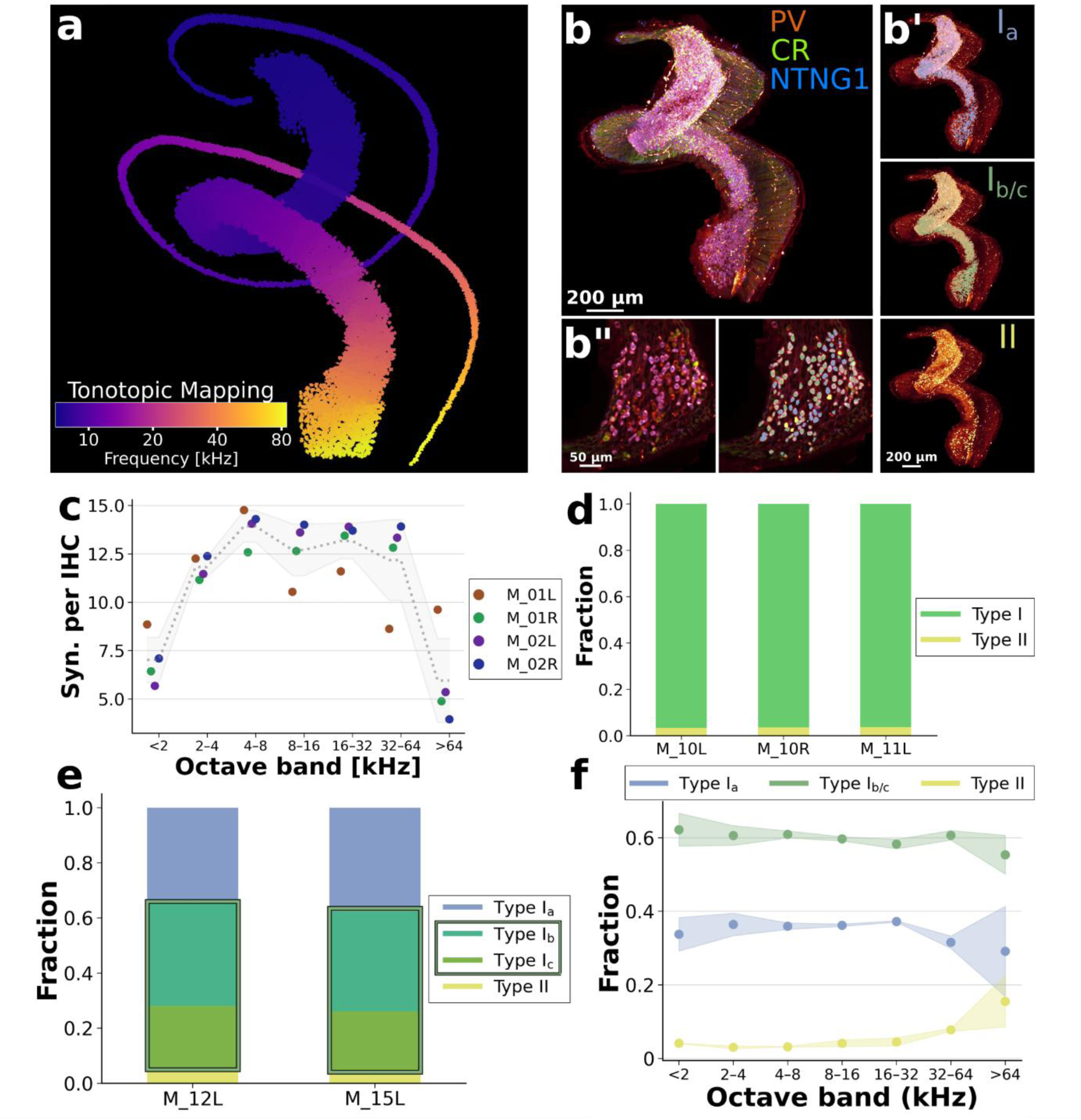
CochleaNet provides tonotopic analysis of the cochlea and SGN (sub)type quantification. **a**, Tonotopic mapping of IHCs and SGNs with frequency bands mapped according to pseudo-greenwood function (Ou *et al*, 2000). **b**, Cochlea with fluorescent SGN (sub)type-specific labels, including rendering of positive expression masks for types I_a_, I_b/c_ and type II SGNs in **b’**. **b’’** shows a zoom-in of fluorescence signal and corresponding (sub)type assignments. **c**, Average ribbon synapse counts per IHC analyzed across frequency bands of four cochleae. **d**, Fraction of type I and type II SGNs identified based on Peripherin (Prph) labelling in three cochleae. **e,** Fraction of subtype I_a_, I_b_, I_c_, and type II SGNs identified based on Ntng1 and Calretinin (CR) labelling in two cochleae. Frame indicates the I_b/c_ fraction. Type II SGNs were identified by negative Ntng1 and CR staining, with result matching type I / type II fractions in **d**. **f**, Tonotopic mapping of subtype fractions from **e**.

Next, we analyzed the representation of type I and type II SGNs as well as type I SGN subtypes along the tonotopic axis. First, we turned to differentiating type II SGNs based on their selective expression of peripherin (Prph, (Hafidi, 1998)). To this end, we prepared three cochleae with PV and Prph staining and segmented SGNs based on PV with CochleaNet. We then measured the median PV and Prph immunofluorescence intensity for all SGN somata and computed the Prph to PV intensity ratio. We manually determined thresholds for positive Prph expression based on this ratio for six sub-volumes along the Rosenthal canal and classified Prph expression for each SGN by applying the threshold of the closest sub-volume. We found a consistent fraction of ca. 0.965 type I SGNs and 0.035 type II SGNs across the three cochleae (Figure 3d, Methods) and an uniform tonotopic distribution (Ext. Data Figure 3b) consistent with literature (Li *et al*, 2020). We then analyzed the location of type II SGNs across the cross-section of the Rosenthal canal, finding that they are preferentially located at the periphery, as expected based on prior work (Hafidi, 1998; Huang *et al*, 2007) (Ext. Data Figure 3c).

We then stained for molecular markers of SGN subtypes I_a_ and I_b/c_ (Petitpré *et al*, 2018; Shrestha *et al*, 2018; Wang *et al*, 2023; Sun *et al*, 2018). Here we used PV as a general SGN marker, calretinin (CR), another Ca^2+^ binding protein that is preferentially expressed in subtype I_a_, weakly expressed in I_b_, and not in I_c_, and for *Netrin-G1(Ntng1)-Cre-*dependent tdTomato expression, which genetically labels subtypes I_b/c_ (Franco *et al*, 2025) (Figure 3b, Ext. Data Figure 3d and e). We proceeded as described for Prph above to identify CR- and Ntng1-expressing cells and classified CR-positive and Ntng1-negative ones as I_a_ and Ntng1-positive ones as I_b/c_. We further divided Ntng1-positive SGNs into CR-positive (putative type I_b_) and CR-negative (putative I_c_). Cells expressing neither CR nor Ntng1 were assigned to SGN type II, as the combined CR/Ntng1 staining exhaustively labels type I SGNs. We found the classifications to be very consistent across cochleae, with average fractions of approximately 0.35 I_a_, 0.60 I_b/c_ (0.38 I_b_, 0.22 I_c_), and 0.05 type II (Figure 3e). Note that the fraction of type II SGNs was consistent with that estimated by Prph staining (Figure 3d). These results largely match previously reported representations of SGN types and subtypes (Shrestha *et al*, 2018; Franco *et al*, 2025; Wang *et al*, 2023; Petitpré *et al*, 2018). We also analyzed the tonotopic distribution of SGN subtypes I_a-c_ and type II, finding their abundance to be largely invariant along the tonotopic axis (Figure 3f). This agrees with RNAseq work (Shrestha *et al*, 2018) but contrasts a previous immunofluorescence-based report of an apicobasal gradients of I_a_ and I_b_ abundance and a reverse gradient of I_c_ (Wang *et al*, 2023). We also prepared three cochleae with staining for Calbindin1 (Calb1), which is expressed in type I_b_ but not I_c_ SGNs, and Ntng1, in addition to PV. We analyzed these cochleae following the same logic as before, finding a fraction of ca. 0.65 I_b/c_ SGNs and fractions of I_b_ SGNs ranging from 0.41 to 0.50 and fractions of I_c_ ranging from 0.18 to 0.25 (Ext. Data Figure 3f and g), overall consistent with our I_b/c_ analysis based on CR and Ntng1, but showing some inconsistencies in the exact I_b_/I_c_ fractions, highlighting the need for further investigation into the exact distinction of I_b_ and I_c_ subtypes.

Overall, we find that CochleaNet analysis is suitable for tonotopic mapping of key morphological parameters of the cochlea and provides access to SGN subpopulations by classifying segmented SGNs according to established markers.

### CochleaNet enables the reliable preclinical evaluation of optogenetic therapies in the cochlea

Future optogenetic hearing restoration will use optical cochlear implants (oCI) to achieve higher spectral selectivity than currently achievable with electrical cochlear implants (eCI) (Huet *et al*, 2024; Wolf *et al*, 2022; Kleinlogel *et al*, 2020). This approach is currently under development and promises improved hearing restoration with more natural perception and improved speech understanding in daily situations with background noise. Here, we tested the utility of CochleaNet for the preclinical evaluation of optogenetic therapies. We immunolabelled and optically cleared ten cochleae of five mice that had been postnatally injected with AAV carrying the new channelrhodopsin variant ChReef (Alekseev *et al*, 2025) under the control of a human synapsin promoter. The green light-activated ChReef mediates large stationary photocurrents and enables sub-microjoule optical stimulation of SGNs in mice and gerbils (Alekseev *et al*, 2025). In order to assess efficiency and specificity of ChReef expression, the treated left and untreated right cochleae were explanted 9-17 weeks after AAV injection, processed, stained for PV, as SGN marker, and GFP, labelling the EYFP-tagged ChReef, and imaged by our high-resolution isotropic light sheet microscope (Figure 4a and b, Methods).

**Figure 4:**
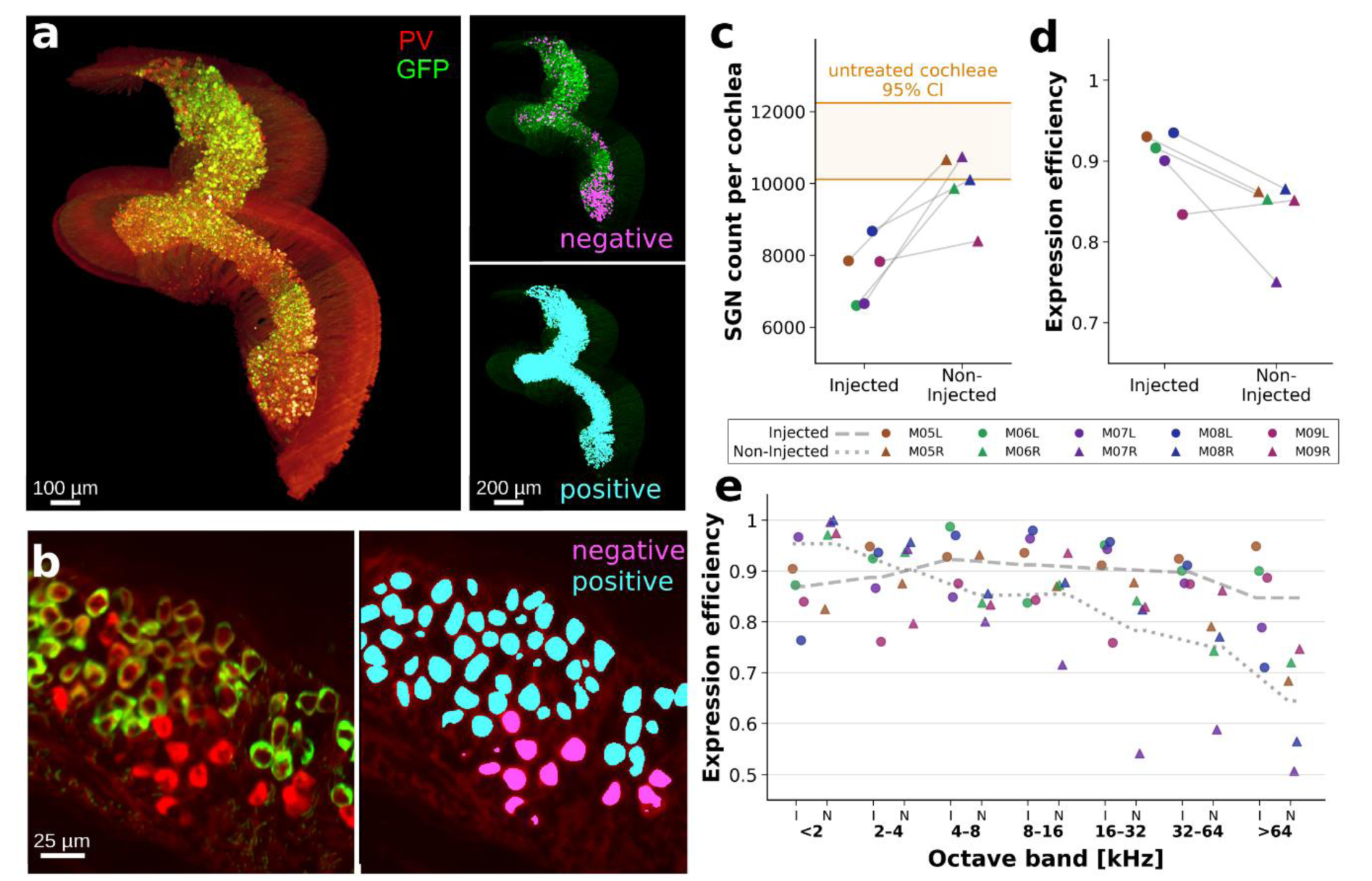
CochleaNet facilitates the evaluation of an optogenetic therapy targeting SGNs. **a**, Cochlea treated with ChReef AAV therapy. Composite of PV and GFP intensities (left) as well as GFP intensities with overlays of GFP-negative (right, top) and GFP-positive (right, bottom) SGNs. **b**, Zoom-in of PV and GFP signals in SGNs (left) with overlay of positive and negative masks (right). **c**, Monitoring of SGN loss in the injected (left) and non-injected (right) cochleae of five animals (M05-M09) treated with the therapy. **d**, Evaluation of expression efficiency determined by analysis of GFP intensity in SGN somata. **e**, Tonotopic mapping of expression efficiencies.

Using CochleaNet, we segmented SGNs based on PV-immunofluorescence and estimated their number as before, here to probe for potential therapy-related, safety-relevant SGN loss (Figure 4c). We found the SGN counts of the injected left cochleae (7,523±791, n=5), and to a lesser extent of the non-injected right cochleae (9,954±846, n=5), to be lower than those of cochleae of untreated mice (11,176±542, n=4, cf. Figure 2d). Substantial ChReef expression was also found in SGNs of the non-injected ear, likely due to viral spread via the cerebrospinal fluid space (Keppeler *et al*, 2018; Kho *et al*, 2000). Therefore, the finding might indicate partial SGN loss due to postnatal injection, AAV exposure, ChReef and/or EYFP expression.

Next, we analyzed the efficiency of the AAV-mediated ChReef expression in SGNs based on the GFP immunofluorescence of SGN somata (Figure 4a and b). To this end, we measured the median GFP immunofluorescence intensity per SGN soma and empirically determined an intensity threshold for successful transduction similar to previous approaches (Huet *et al*, 2021). Due to the inhomogeneous immunofluorescence intensities across the cochlear volume, we determined thresholds for six sub-volumes extracted uniformly along the tonotopic axis of each cochlea. For each SGN, we then applied the threshold of the closest sub-volume. ChReef expression efficiency was high for both the injected left cochleae (0.903±0.037, n=5) and the non-injected right cochleae (0.836±0.043, n=5) (Figure 4d). We also analyzed the ChReef expression efficiency along the tonotopic axis for injected and non-injected cochleae (Figure 4e). Consistent with literature (*e.g.* Keppeler *et al*, 2018), we found a trend toward an apicobasal gradient of ChReef expression efficiency that was more pronounced in the non-injected cochleae.

In summary, we conclude that CochleaNet combined with LSFM is a powerful tool for assessing key preclinical efficiency and safety data for optogenetic therapy of the cochlea.

### CochleaNet generalizes to different stainings, imaging conditions and animal species

Next, we tested the versatility of CochleaNet by transferring sample preparation, imaging, and analysis pipeline to the cochleae of gerbils (Meriones unguiculatus). Given their “human-like” low frequency hearing and larger cochlea (∼2.5 times smaller than human compared to ∼5.0 in mouse, *e.g.* (Keppeler *et al*, 2021)), this rodent species is a popular animal model in translational hearing research (Castaño-González *et al*, 2024; Dieter *et al*, 2020). Here, we stained one cochlea for PV, Vglut3, and CtBP2. Imaging was performed with the high isotropic resolution LSFM (Aakhte *et al*, 2025). We segmented SGNs and IHCs, and detected synapses, as described before. Here, the CochleaNet networks for SGN and ribbon synapse analysis yielded high-quality results – judged by manual inspection – as is. For IHCs, the segmentation quality suffered from the different intensity profile of the Vglut3 signal in gerbil IHCs. Hence, we annotated 129 cells in 9 additional sub-volumes and retrained the network on this data to improve IHC segmentation quality (Figure 5a and b). We estimated 18,541 SGNs, 1,180 IHCs, and 18.3±8.5 ribbon synapses per IHC in this exemplary cochlea and compared to previously reported values (Figure 5c, Ext. Data Table 3).

**Figure 5:**
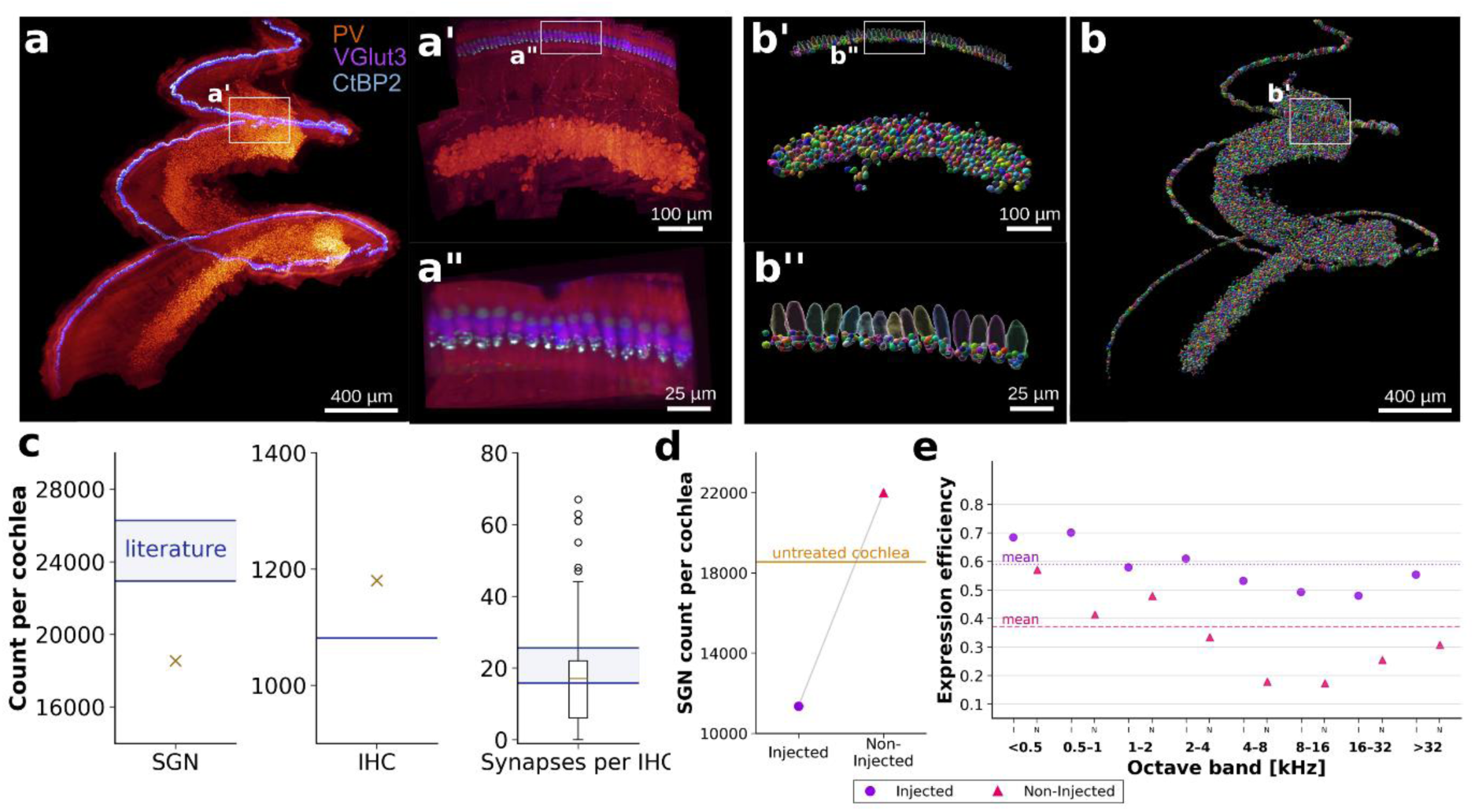
CochleaNet generalizes to analyses of the gerbil cochlea. **a**, Gerbil cochlea imaged in the high-resolution microscope with PV (red), Vglut3 (blue) and CtBP2 (white) staining; right panels show the respective signals at higher magnification. **b**, SGN and IHC segmentation as well as synapse detection (small spheres in zoom-in) from CochleaNet. **c**, Quantification of gerbil SGN, IHC, and ribbon synapse counts with literature values. **d**, SGN counts of a gerbil treated with f-Chrimson therapy, for injected (left) and non-injected (right) cochlea, with reference value from the untreated animal. See Ext. Data Figure 4c for the corresponding SGN densities. **e**, Efficiency of f-Chrimson expression derived from GFP intensities, analyzed across the tonotopic axis.

Another gerbil received a postnatal optogenetic therapy for expressing the fast-closing red light-activated channelrhodopsin variant f-Chrimson in SGNs (Mager *et al*, 2018; Huet *et al*, 2021). We labelled the injected left cochlea and non-injected right cochlea for PV and GFP, the latter marking f-Chrimson-EYFP expression, respectively, and segmented the SGNs in these cochleae (Ext. Data Figure 4a and b). First, we compared the SGN counts, finding 11,351 somata in the left injected and 21,995 somata in the right non-injected cochlea (Figure 5d), compared to 18,541 in the untreated animal. Note that part of the Rosenthal canal in the left injected cochlea as well as the untreated cochlea were torn off during preparation, partially explaining the differences in cell counts. For further analysis, we measured the SGN density along the Rosenthal canal, finding very similar values in the cochlea of the untreated gerbil and the non-injected cochlea, but a lower density in the injected cochlea (Ext. Data Figure 4c), indicating SGN loss related to the AAV-f-Chrimson therapy. Next, we measured GFP immunofluorescence intensity per SGN soma and determined f-Chrimson expression based on intensity thresholds determined manually for eleven (injected cochlea) and nine (non-injected cochlea) sub-volumes along the Rosenthal canal. We found an expression efficiency of 0.589 in the injected and 0.379 in the non-injected cochlea which were rather uniform along the tonotopic axis (Figure 5e).

Finally, we tested CochleaNet’s utility to analyze data from commercially available LSFMs. First, we turned to data acquired by (Keppeler *et al*, 2021) on two mouse cochleae and one marmoset cochlea, each labelled for PV and myosin VIIA (Myo7a). Myo7a is an unconventional motor protein specifically expressed in cochlear IHCs and outer hair cells. These samples were imaged with an UltraMicroscope II (Miltenyi Biotec). Due to the low and anisotropic resolution of this imaging configuration (∼1µm x 1µm x 5µm), we trained new versions of the CochleaNet networks for SGN and IHC segmentation based on newly annotated training data for SGNs (18 sub-volumes of the PV-signal with 1,416 cells) and IHCs (17 sub-volumes of the Myo7a signal with 610 cells). These networks faithfully segmented SGNs and IHCs as judged by manual inspection. We found 9,000 and 9,434 SGNs as well as 583 and 544 IHCs in the two mouse cochleae (Figure 6a and d) and 21,253 SGNs and 1,285 IHCs in the marmoset cochlea (Figure 6d, Ext. Data Figure 5). These values closely matched the SGN counts determined for the mouse cochleae by (Keppeler *et al*, 2021) but are lower compared to the high-resolution LSFM data (cf. Figure 2). We hypothesize that this is because closely packed cells cannot always be distinguished, due to the low resolution and anisotropy of the data. To validate this hypothesis, we compared the average volume of SGNs and IHCs (Ext. Data Fig 5b) between segmentations of the high-resolution microscope (samples from Figure 2) and the low-resolution microscope. The results show a larger average volume and more outliers with larger volume in the samples from low-resolution LSFM, validating our hypothesis. Also note that (Keppeler *et al*, 2021) did not quantify IHCs in the mouse cochleae or SGNs and IHCs in the marmoset cochlea.

**Figure 6:**
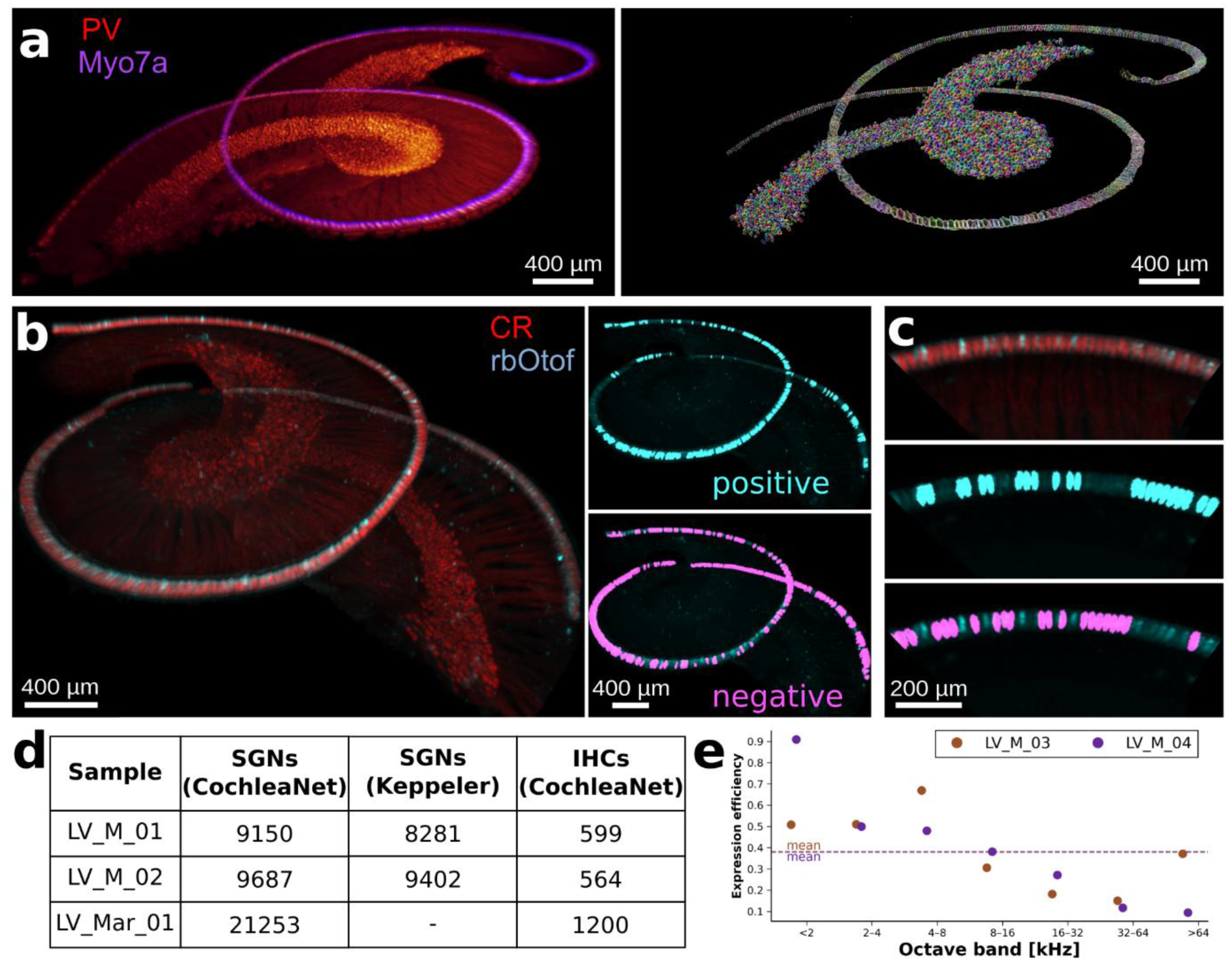
CochleaNet extends to the analysis of lower-resolution LSFM data from a commercially available microscope. **a**, Mouse cochlea with PV (red) and Myo7a (blue) labelling (left) and derived SGN and IHC segmentation (right). Data from (Keppeler *et al*, 2021). **b**, Mouse cochlea with OTOF gene replacement therapy (Rankovic *et al*, 2021), with CR (red) and rbOTOF (cyan) labelling (left). IHCs were segmented based on CR and classified into otoferlin-expressing (top right) and non-expressing (bottom right) cells based on the rbOTOF signal. **c**, Zoom in of CR and rbOTOF signal in IHCs (top) as well as corresponding positive (middle) and negative (bottom) expression masks, overlayed with rbOTOF signal. **d**, Quantification of SGN and IHC counts in the samples from (Keppeler *et al*, 2021) with comparison to their previously determined values (only measured for SGNs in mice). **e**, Tonotopic mapping of rbOTOF expression efficiency in two cochleae from (Rankovic *et al*, 2021).

For another exemplary cochlear data set acquired with an UltraMicroscope II, we turned to a previous study preclinically characterizing *OTOF* gene replacement therapy in *Otof^-/-^* mice that targets IHCs. Pathogenic mutations in *OTOF* disrupt synaptic transmission at the ribbon synapse of IHCs, in which otoferlin plays a central role (Moser & Starr, 2016). *OTOF* gene therapy has entered clinical trials as the first inner ear gene therapy with promising safety and efficiency data (e.g. Qi *et al*, 2024; Lv *et al*, 2024). Given the challenges, including the large size of *OTOF* and the preliminary nature of the preclinical and clinical efficacy data, further preclinical and clinical studies are warranted. Here, we used CochleaNet to evaluate the previously described overload-AAV approach (Rankovic *et al*, 2021), for a preliminary analysis of *OTOF* expression in an entire *Otof^-/-^* cochlea (Figure 6b and c). We took advantage of data acquired 18 days after cochlear injection of PHP.eB-CMV/HBA-fl-Otof on postnatal day 6 with a commercial LSFM (UltraMicroscope II; (Rankovic *et al*, 2021)). The IHCs in this data were stained with calretinin. We segmented them with CochleaNet using the network trained on low-resolution Myo7a signal (see previous paragraph) and found that it faithfully segmented IHCs despite the different staining. When comparing counts of IHCs to those of healthy and untreated mice (cf. Figure 6d), we found no indication of IHC loss, estimating 643 and 552 IHCs respectively, compared to 582±18 IHCs in the healthy animals imaged with the same microscope. Next, we evaluated the otoferlin expression efficiency. Here, we had to take a different approach compared to optogenetic therapies targeting SGNs (cf. Figures 4 and 5) due to a slight misalignment of the CR signal and the rbOTOF signal labelling AAV-induced gene expression. To alleviate this misalignment, we dilated the segmented IHC masks and calculated different statistics of the rbOTOF signal aggregated over dilated masks, which were then used as features for a random forest classifier trained on manually identified examples of positively and negatively expressing cells (Methods). We applied this classifier to all IHC masks and visually validated the results, finding an overall accurate classification with some inconsistencies in intensities that were assigned to multiple dilated cells. We then analyzed the expression efficiency for both cochleae, finding otoferlin expression in a fraction of 0.38 of IHCs, and analyzed the expression along the tonotopic axis (Figure 6e). We also compared the CochleaNet-based approach to the previous manual analysis (Rankovic *et al*, 2021), finding that we measured slightly higher overall expression efficiencies, likely due to including more IHCs in our analysis (Ext. Data Figure 5d).

In summary, this section demonstrates that the utility of CochleaNet extends beyond reproducible segmentation of high isotropic resolution LSFM data of the mouse cochlea. CochleaNet generalizes to analyses of mouse cochlear data obtained with different stainings and imaging conditions as well as to data from the larger cochleae of gerbils and marmosets.

## Discussion

We have developed CochleaNet, a comprehensive framework for the analysis of LSFM data of the cochlea. CochleaNet accurately segments two key cochlear cell types: SGNs and IHCs, and faithfully detects their ribbon synapses. We applied CochleaNet to a large set of mouse cochleae to analyze their morphology in detail and to evaluate optogenetic and gene replacement therapies targeting SGNs and IHCs, respectively. Finally, we demonstrated CochleaNet’s versatility for application to data acquired from the mouse cochlea with different stainings or microscopes and data from the larger cochleae of gerbil and marmoset. We expect CochleaeNet to automate the challenging analysis of cochlear LSFM data, thereby facilitating the broader use of LSFM as a standardized, powerful analysis tool. As illustrated, the combination of advanced LSFM and CochleaNet can serve the evaluation of novel therapeutic approaches of the cochlea including gene and optogenetic therapies as well as new drugs (Rommelspacher *et al*, 2024) and cell therapies (Chen *et al*, 2012). Moreover, the utility of CochleaeNet to analyze cochleae of other species will enable work with other relevant animal models such as other rodents and non-human primates as well as with postmortem human cochleae, with no or little training on additional annotated samples.

### Advancing the analysis of whole-organ fluorescence microscopy imaging

General-purpose pretrained models (Stringer *et al*, 2021; Schmidt *et al*, 2018; Archit *et al*, 2025) continue to struggle with efficient and reliable segmentation/detection of cells and subcellular structures in large volumes such as in LSFM data of the whole cochlea <u>(Keppeler et al. 2021)</u>. While more targeted approaches have been developed (Buswinka *et al*, 2023b; Cai *et al*, 2025), a solution for the analysis of different cochlear cell types and subcellular structures, such as synapses, had been missing. In particular, the reliable quantification of the afferent neural connectome of the normal, diseased and treated cochlea with its intricate spiral-shaped organization remained an important task. By advancing deep learning-based analysis of high quality cochlear LSFM data, CochleaNet achieved the reliable analysis of IHCs, SGNs and synaptic ribbons at the IHC-SGN synapses in the intact rodent cochlea. Important future tasks *en route* to elucidating cochlear afferent and efferent connectivity include segmentation of outer hair cells as well as afferent and efferent neurites and their synapses. Domain adaptation and foundation models will be key for such further developments. We foresee that additional annotation efforts will also be required for neurites, which appear as thin fibers in the data (*e.g.* Figure 1, see also (Aakhte *et al*, 2025)). Yet, compared to the complexity of brain connectomes targeted in recent expansion microscopy work (Tavakoli *et al*, 2025), the cochlea’s spatial organization and relatively small size promise feasible solutions for the entire organ. In addition, CochleaNet demonstrated versatility to cope with other stainings, imaging conditions and species. Eventually, the approach illustrated for the cochlea is more generally applicable to analyze the multicellular organization of tissues and organs such as the retina or the whole eye as well as heart muscle or the whole heart.

### CochleaNet – a framework for analyzing the molecular anatomy of the cochlea in the normal, diseased and treated state

Combining LSFM and CochleaNet fills an important gap in analyzing the molecular anatomy of the cochlea in the normal, diseased, and treated state. Analysis of the afferent connectome, has largely relied on EM, fluorescence microscopy, and X-ray microscopy. The spatial scales covered by these approaches range substantially, overlap partially, and each approach has complementary benefits and limitations. For example, EM ranges from the molecular to the tissue scale, while photonic imaging ranges from the organ to the subcellular scale. EM has been key for the analysis of afferent synapses (Smith & Sjöstrand, 1961; Liberman, 1980; Michanski *et al*, 2019; Hua *et al*, 2021). However, fluorescence imaging in combination with knock-out validated molecular synaptic markers (Khimich *et al*, 2005; Jean *et al*, 2018; Roux *et al*, 2006) readily offers molecular information and a larger spatial scope. This, together with ease of technique and broad availability of confocal microscopes, has made juxtaposed immunofluorescence spots for presynaptic CtBP2/RIBEYE and postsynaptic density (Khimich *et al*, 2005) the arguably most commonly used definition of afferent synapses in health and disease (Kujawa & Liberman, 2019; Moser & Starr, 2016). Yet, unlike X-ray imaging, both approaches typically cover only fractions of the cochlea, while analysis of the entire organ is relevant given differences of hair cells, SGNs, and synapses along the tonotopic axis (Meyer & Moser, 2010; Fettiplace, 2023). This also extends to common cochlear pathologies, such as noise-induced and age-related hearing loss, which show a tonotopic gradient with stronger alterations prevailing in the high frequency base (Aryal *et al*, 2025). For example, loss of synapses and SGNs due to loud noise exposure or aging is most prominent in the cochlear base (Kujawa & Liberman, 2009; Sergeyenko *et al*, 2013). LSFM and CochleaNet will serve future studies that comprehensively analyze changes in number and properties of SGNs, IHCs, and their synapses in response to noise exposure, drugs, or aging as a function of tonotopic position. Future extension of the CochleaNet framework to segment further cell types will extend these analyses to additional cochlear structures and possibly to the vestibular system.

Furthermore, in combination with tissue expansion, LSFM promises to resolve protein clusters in the intact cochlea. Together with future deep learning-based image analysis within the CochleaNet framework, this might even provide low resolution structures of protein complexes as recently shown for confocal imaging of expanded neural samples (Shaib *et al*, 2024). Moreover, the more molecular information we gain on these cells and their subcellular specializations, the more relevant will be molecular markers, multiplexed imaging, and efficient analysis of the entire cochlea, as feasible by combining LSFM and CochleaNet. For example, CochleaNet robustly identifies type I SGNs and their subtypes (Shrestha *et al*, 2018; Sun *et al*, 2018; Petitpré *et al*, 2018). Their function and afferent connectivity along the tonotopic axis remain an important target for molecular anatomical studies. This will not be limited to immunolabeling but can also be extended to other labels, e.g. of RNA. Elucidating the molecular anatomy and pathology of the afferent connectome is key to understanding auditory synaptopathy and neuropathy. Moreover, as exemplified, LSFM imaging and CochleaNet will facilitate the development and evaluation of precision medicine approaches such as gene therapy and optogenetic therapies of the cochlea. The utility of CochleaNet to quantify IHCs and SGNs, which are the targets of *OTOF* gene therapy and optogenetic therapy, respectively, makes it a premier tool to assess efficiency, specificity and longevity of these AAV-based therapies in preclinical studies. For example, demonstrating the absence of channelrhodopsins from IHCs in optogenetically modified but otherwise intact cochleae or the absence of IHCs following ototoxic deafening is highly relevant for assessing specificity and efficiency of optogenetic hearing restoration. Likewise, evaluating *OTOF* gene therapy aside from quantifying transduction efficiency, specificity and expression levels in IHCs as well as the synaptic and neural status of the cochlea, which are known to be compromised by loss of otoferlin expression (Roux *et al*, 2006).

We envision that a similar combination of LSFM imaging and deep learning-based image analysis, as exemplified here for the cochlea, will also support the preclinical evaluation of gene and optogenetic therapies of other sensory organs and systems, such as the retina, the eye, or the vestibular system.

### Limitations

A limitation of the current study arose from the requirement for extraction, fixation and processing of the cochlea, prohibiting functional studies which is shared with most if not all histological approaches. Likewise, relying on immunolabeling, with potential non-linearities of the imaging process and trial-to-trial variability, even with identical protocol and conditions, challenges the quantification of protein levels. We are still far from analyzing all relevant cell types of the cochlea, which would not only require further networks to be trained but also multiplexing of labelling and imaging. The latter could use approaches such as DNA-paint (van Wee *et al*, 2021) and further developed versions of CochleaNet might assist such analysis by general segmentation of all cells and subsequent filtering of the specific cell types. Moreover, the analysis of fluorescent markers labelling gene therapy products was limited to the median expression over SGN and IHC somata. However, the localization of the transgenic protein within the soma can be critical for certain therapies, in particular for optogenetic treatments which rely on channelrhodopsin expression at the cell membrane. The high-resolution LSFM data and precise segmentation masks should enable such analyses in the future. In addition, with only one cochlea of an untreated gerbil, further work is required to provide reliable statistical estimates of SGN, IHC and synapse counts of this species. Furthermore, the analysis of lower resolution data from commercially labelled LSFM is limited to counting and spatial analysis of cochlear cells. It does not enable a detailed analysis of cellular morphology, is prone to underestimating the count of closely packed cells, and does not provide ribbon synapse detections.

## Acknowledgments

This work was supported by the German Research Foundation (Deutsche Forschungsgemeinschaft, DFG) through grants to T.M., J.H. and C.P. under Germany’s Excellence Strategy—EXC 2067/1-390729940 and SFB1690/A04, B01 (to T.M.), SFB1690/A04 (to K.K.). The work was further funded by the Else Kröner Fresenius Foundation through the Else Kröner Fresenius Center for Optogenetic Therapies to K.K. and T.M.. C.P. acknowledges the support of the Ministry of Science and Culture of Lower Saxony through funds for the CAIMed project (grant no. ZN4257). T.M. acknowledges the support by the European Research Council through the Advanced Grant “DynaHear” under the European Union’s Horizon 2020 Research and Innovation program (Grant Agreement No. 101054467). M.A. and J.H. acknowledge funding by the Alexander von Humboldt Foundation (AvH professorship to J.H.). We also gratefully acknowledge the computing time granted by the Resource Allocation Board and provided on the supercomputer Lise and Emmy at NHR@ZIB and NHR@Göttingen as part of the NHR infrastructure. The calculations for this research were conducted with computing resources under the project nim00007. The work was further funded by Fondation Pour l’Audition (FPA RD-2020-10) and the Volkswagen-Stiftung from the “Niedersächsisches Vorab” (ZN3898 and ZN4000) to T.M.. LR was supported as clinician scientist by EXC2067 and the Else Kröner Fresenius Center for Optogenetic Therapies.

We thank the following people: Sarah Muth and Julia Jeremias for providing an implementation of the network used for synapse detection. Victoria Hunniford and Anna Vavakou for providing ChReef treated cochleae. Niels Albrecht for providing gerbil cochleae and Nina Rothenberg for aiding in the clearing of G_1L and G_1R. Gerhard Hoch, Ina Preuß and Sandra Gerke for expert technical support. Patricia Räke-Kügler for expert administrative support. Jakob Neef for expert support in data storage, data handling and further technical assistance. Alexey Alekseev for advice on data evaluation. Bettina Wolf for scientific discussions and early supervision. Nare Karagulyan for discussions on ribbon synapse annotation. Anwai Archit for feedback on figures.

## Author Contributions

L.R, A.M.D., K.K., J.H., T.M., and C.P. conceptualized the work. M.S. and C.P. implemented the software. L.R., A.M.D., E.K., M.U., and A.T. performed sample preparation, image acquisition, or data annotation. M.A. built and optimized the isotropic light-sheet microscope. J.H. provided resources for the light-sheet experiments. L.R., A.M.D., E.K., M.S., and C.P. analyzed data. L.R., M.S., M.A., and C.P. visualized the results. L.R., A.M.D., T.M. and C.P. drafted the manuscript. All co-authors reviewed and edited the final manuscript.

## Methods

### Animals

All animal experiments were carried out in accordance with relevant national and international guidelines (European Guideline for animal experiments 2010/63/EU, German Animal Welfare Act). The procedures have been approved by the responsible office of the regional government. Rodents were kept in a 12 h light/dark cycle with ad libitum access to food and water as well as enrichment. Here, data was obtained from C57BL/6 wild-type mice, NetrinG1-Cre/ai14 reporter mice (Bolding *et al*, 2020), Mongolian gerbils (Meriones unguiculatus) and *Otof*^−/−^ mice (Reisinger *et al*, 2011) of either sex between the ages of 4 and 52 weeks. An overview of all specimen used in this study can be found in Ext. Data Table 1.

### Gene therapy in mice and gerbils

Viral vector purification procedure of ChReef was performed as previously published, and detailed descriptions of its process are available in (Huet & Rankovic, 2021). The final constructs used for gene therapy in rodents consisted of either ChReef (Alekseev *et al*, 2025) for mice or f-Chrimson (Mager *et al*, 2018) for gerbils, each under control of the human synapsin promoter (neuronal targeting) and tagged with the reporter eYFP for histological analyses. While ChReef was packed into AAV2/9 together with an enhancement of the trafficking signal and export signal, f-Chrimson was packed into PHP.B.

Postnatal cochlear administration of the gene therapy was performed for all animals used in this study as previously published (Keppeler *et al*, 2018; Mager *et al*, 2018; Huet & Rankovic, 2021; Alekseev *et al*, 2025). Briefly, rodents were isoflurane-anesthetized (5% for anesthesia induction, 1–3% for maintenance with frequent monitoring of the hind-limb withdrawal reflex and corresponding anesthesia adjustments) and surgeries to administer corresponding gene therapies were performed under general analgesia (local xylocaine as well as buprenorphine (0.1 mg/kg) and carprofen (5 mg/kg) administration). Further, physiological body temperature was maintained throughout surgeries. Here, each animal was injected through a small retroauricular incision spreading the tissue surrounding the respective cochlea carefully to expose the round window. Then, the round window membrane was gently punctured and the viral vector solution was administered with a quartz or borosilicate capillary pipette at a postnatal age of 6 days for mice and seven days for gerbils. Here, the following viral constructs and titers were injected:

– Single AAV optogene therapy:

- AAV2/9-hsyn-ChReef-TS-EYFP-ES at 2.45E + 13 GC/ml (cdPCR) to C57BL/6 wild-type mice
- PHP.B-hSyn-fChrimson-EYFP at 1.31E +13 GC/ml (cdPCR) to wild-type gerbils

Following the injection, the tissue was repositioned and the wound was sutured. For postoperative analgesia Carprofen (5 mg/kg) was administered for one day the day after surgery. At 9-19 weeks after injection mice and gerbils were euthanized, the cochleae were extracted and further processed.

### Sample preparation

After extraction from the temporal bone, cochleae were fixed for 45 minutes for mice and 60 minutes for gerbils in 4% formaldehyde and subsequently rinsed in phosphate-buffered saline (PBS). Samples were then decalcified for four to seven days week in 10% ethylenediaminetetraacetic acid (pH 8). After decalcification, cochleae were extensively trimmed with surgical instruments (Fine Science Tools) to remove as much bone as possible for easy access and sufficient penetration of the light sheet beam during optical sectioning. For reproducible immunolabelling as well as optical clearing, we used a newly optimized iDISCO^+^ clearing protocol (adapted cDisco^+^ from (Keppeler *et al*, 2021)). Here, we decolorized the decalcified cochleae for two days using 25% amino alcohol (N,N,Nʹ,Nʹ-Tetrakis(2-Hydroxypropyl)ethylenediamine). Next, we employed pretreatment and permeabilization of the samples for approximately four days according to iDisco (Renier *et al*, 2014), specifically without the use of methanol, followed by epitope blocking. For reliable immunolabelling of the entire organ including IHCs, SGNs and subcellular structures, sufficient penetration of the cochleae was achieved by incubating each antibody for seven to 14 days for mice and gerbil cochleae, each followed by washing in PBS-Tween 0.2% with Heparin 10μg/mL for up to one day (Duque Afonso, 2020; Keppeler *et al*, 2021). All antibodies and their combinations used in this study can be found in a respective table attributed to each sample and condition (Ext. Data Table 1 and 2). Following the immunolabelling, the cochleae were dehydrated in a stepwise manner using an ethanol or methanol/H_2_O series ranging from 20 to 100%. Subsequently samples were delipidized with dichloromethane (DCM, (Renier *et al*, 2016)). Finally, ethyl cinnamate (ECi) was used to match a RI of 1.559 and was also present for sample storage and imaging.

### Image acquisition and data preprocessing

For acquiring the high-resolution LSFM volumes, we have used the recently developed high-speed, aberration-corrected light sheet fluorescence microscope, which was optimized for isotropic resolution of cleared tissue at sub-micron resolution (Aakhte *et al*, 2025). In brief, our setup achieves an isotropic resolution of 850 nm in both axial and lateral dimensions, enabling accurate recording of structural features across large cochlear volumes. In the detection arm of the microscope, we have employed a 16X multi-immersion dipping objective (ASI 15-10-12, NA 0.4, WD ∼12 mm) to cover high refractive index (RI) solvents up to 1.56. Further, the detection lens was paired with a 200 mm tube lens (TTL200-A, Thorlabs), projecting an image onto a scientific complementary metal-oxide-semiconductor camera (PCO Panda 4.2m) with 6.5 µm pixel pitch. At this magnification and RI, the effective pixel size at the sample plane was 380 nm, which matches the Nyquist criterion for diffraction-limited resolution. In the illumination arm, the laser source includes a free-beam laser engine with outputs at 405, 488, 561, and 640 nm with a maximum laser power of 100 mW for each line. The illumination objective was a 20X air objective (Mitutoyo Plan Apo, NA 0.42, WD 20 mm) positioned outside the sealed sample chamber to eliminate the need for solvent-compatible immersion. To additionally minimize spherical aberrations arising at the air-glass-medium interface, we introduced a positive meniscus lens (f = 100 mm, Thorlabs LE1234) between the illumination lens and the sample chamber. The sample chamber was designed with five openings: one for dipping the sample from the top, one for holding the detection objective lens, and one for holding the meniscus lens. The other two windows were used for LED-based preview illumination and for visual access to the cochlea during tiling adjustments. The chamber was fabricated from anodized aluminium and sealed with chemically resistant O-rings compatible with ethyl cinnamate (ECi).

To mount the RI–matched cochlea a customized micro-clamp was used to securely hold the cochlea in a 3D-printed holder with a simple connection to a motorized stage, allowing movement in four dimensions (XYZR stage). The R stage helps determine the optimal orientation of the cochlea prior to 3D imaging. Since the detection optical arm’s field is about 800 μm x 800 μm, we performed mechanical tiling using X and Y stages to image the entire cochlea. The Z stage enables fast and isotropic volumetric imaging by continuously translating the cochlea through the light sheet with precise Z-plane spacing down to 380 nm at 40 frames per second per plane. To keep the imaging smooth and as fast as possible, all volume tiles were imaged for each wavelength separately, meaning the first volume tile was captured, followed by the second, and so on, for each excitation wavelength. Simultaneously with recording, each acquired image stack was streamed directly to an SSD (Samsung SSD, 4 TB) via Thunderbolt 4, with a writing speed of 1 Gbit per second, for minimal data latency. Data was saved in RAW format in real-time during imaging.

After completing multi-tile volume imaging, the RAW data were converted to n5 file format using functionality that is part of the Python library we developed. These files were loaded in BigStitcher (Hörl *et al*, 2019), which was used for stitching the volume tiles to each other and to align the different channels. We first realigned tiles based on the imaging sequence with the *Move Tiles to regular Grid* option under *Arrange Views* in the *Preprocessing* tab. We then used the *Stitching Wizard* for initial alignment of tiles and channels. We further enhanced the tile and channel alignment using *Interest Point Stitching* and *Registration Refinement*. After visual confirmation of registration, the volumes were exported to OME-Zarr (Moore *et al*, 2021) and were copied onto S3 storage so that they could be accessed with MoBIE (Pape *et al*, 2023), which enabled seamless browsing of the volumes via web-access.

### Data annotation

Data annotation was performed by three co-authors with expertise in cochlear biology. They annotated SGNs and IHCs as precise annotation masks for network training and with points marking the center for network evaluation. For ribbon synapses, one co-author annotated synapse locations as points, which were used for network training and evaluation.

The SGN segmentation masks for network training were annotated in small sub-volumes of the PV stain by correcting initial segmentation results from CellPose (Stringer *et al*, 2021) with the paint mode of napari. Each volume was annotated by a single expert; experts consulted on difficult cases. In total, 30 sub-volumes were annotated this way. The sub-volumes came from seven different cochleae (IDs: M_18L, M_06L, M_10L, M_19R, M_17R, M_05L, M_20R) and contained a total of 2,782 annotated SGN somata. In addition, we extracted 18 sub-volumes that did not contain any SGN somata, from five different cochleae (IDs: M_18L, M_06L, M_21L, M_17R, M_15R). These sub-volumes showed off-target signal in PV, for example in afferent neurites or in bones. They were included in the model training to suppress unspecific segmentation output. For training the lower-resolution version of the SGN segmentation model, 18 sub-volumes with PV signal extracted from the data of (Keppeler *et al*, 2021) with 1,416 cells were annotated.

The IHC segmentation masks were annotated following a similar procedure as for SGNs. Here, we used an initial segmentation from µSAM (Archit *et al*, 2025), which performed better than CellPose for this task. We annotated a total of eleven sub-volumes from four samples (IDs: M_22L, M_23L, M_24R, M_01L). They contained a total of 283 annotated cells. We added one sub-volume containing no IHCs, but with off-target signal, from sample M_01L. The IHC segmentation model for gerbil data was trained on nine additional sub-volumes extracted from a gerbil cochlea (G_1L) containing 129 annotated IHCs, in addition to the previously described annotated IHC data from mice cochleae. For training the lower-resolution version of the IHC segmentation model, 17 sub-volumes with Myo7a signal extracted from the data of (Keppeler *et al*, 2021) with 610 cells were annotated.

Ribbon synapses were annotated by a single expert using a semi-automated workflow in IMARIS. The *Add new spots* tool was used to detect synapses in a sub-volume containing an IHC row with IHC (Vglut3, Myo7a, or CR) and synapse (CtBP2) staining based on intensity and size thresholds. The automated detection results were then corrected manually. This way, 15 sub-volumes from six different cochlea were annotated (IDs: M_25L, M_26L, M_18L, M_27R, M_26R, M_01R) for training. They contained 3,996 annotated synapses. In addition, we annotated seven sub-volumes from three cochlea (IDs: M_01L, M_02L, M_02R) for network evaluation following the same procedure as described above. They contained 1,644 annotated synapses.

For evaluating the SGN segmentations, we annotated the center positions of the cells visible in twelve complete image sections (PV stain) from five samples (IDs: M_02L, M_28R, M_21L, M_01L, M_02R). The sections were first annotated individually by each expert, with a context of ten sections above and below to judge if the signal in the central section corresponds to a SGN or some other structure. We then programmatically matched these annotations to each other by counting two or more annotations closer than three micrometers as a positive match and declaring other annotations as non-matching. In a second round, the experts then independently reviewed all non-matching annotations from the first stage and marked the ones that, in their estimate, corresponded to a SGN missed. These annotations were matched to each other following the same approach as before and the consensus annotations from the first and second stage were combined to obtain the final consensus annotations. In total, we obtained 2,565 consensus SGN annotations. We chose this procedure rather than using held-out data with annotated segmentation masks to evaluate the segmentation for a broad range of areas in multiple cochleae. Achieving a similar coverage with mask annotations would not have been possible due to the large effort involved in creating mask-level annotations.

For evaluating the IHC segmentations we chose a similar approach as for SGNs with three experts marking the cell centers in twelve image sections (Vglut3) from four samples (IDs: M_01L, M_01R, M_02L, M_02R). In this case, we only performed a single round: the annotations were created independently and then matched to each other to obtain a consensus based on a threshold of six micrometer. The two-stage process used for SGNs was not necessary because annotating IHCs was easier than SGNs due to less background signal, a more regular spatial configuration, and less cells per section. We obtained 320 consensus annotations for IHCs.

### Network architectures, training, prediction, and post-processing

We trained three different networks for the analysis of cochleae in LSFM: for the instance segmentation of SGNs based on PV stain, for the instance segmentation of IHCs based on Vglut3 stain, and for the detection of ribbon synapses based on CtBP2 stain. The networks share the same 3D U-Net architecture (Falk *et al*, 2019) with 32 initial feature channels, four encoder-decoder levels, and the number of features channels increasing by a factor of two in each encoder level. For SGN and IHC segmentation, two distinct versions of the networks were trained for segmentation of high-resolution isotropic data and for segmentation of lower-resolution data.

For SGN and IHC segmentation, we trained the respective networks to output three channels, predicting foreground probabilities, regressing the normalized distance to the cell center, and regressing the normalized distance to the closest cell boundary. We used the negative dice score as loss function for all three outputs, with targets derived from the respective cell mask annotations. For distance outputs, the loss was computed only inside of cells. The networks were trained for 100,000 iterations with an initial learning rate of 1e-4 and a learning rate scheduler that reduced it by a factor of 0.5 if the validation loss plateaued for 5 epochs. We extracted the model checkpoint of the epoch with lowest validation loss, which was computed after each epoch on a separate validation split that made up 6.3% of the data for SGNs and 8.3% for IHCs. During inference, we segmented instances (individual cells) by applying a threshold to the distance outputs to obtain seeds for watershed-based segmentation, using the binarized foreground probabilities as mask, and the boundary distance output as heightmap. Specifically, we used the following parameters for SGNs:

- foreground_threshold = 0.5,
- boundary_distance_threshold = 0.5,
- and center_distance_threshold = 0.4, and the following for IHCs:
- foreground_threshold = 0.5,
- boundary_distance_threshold = 0.6,
- center_distance_threshold = 0.5,
- and distance_smoothing = 0.6.

This segmentation procedure was adopted from µSAM (Archit *et al*, 2025). For application to whole cochleae, we implemented additional post-processing: For SGNs, we computed the centroids of all segmented objects and then built a spatial graph that connected all pairs of cells closer than a given minimal distance with an edge. The minimal distance was set to a default of 30 µm and was increased to up to 70 µm for individual cochleae to account for small gaps within the Rosenthal’s canal. We computed the connected components of this graph and retained only the largest component, which typically covered the SGNs of the Rosenthal canal, while removing segmented objects corresponding to off-target signal. In cases where parts of the Rosenthal canal were torn off during sample preparation, we also included the SGNs from the corresponding components. For IHCs we applied the same approach. In addition, we post-processed the IHCs after matching ribbon synapse detections (next paragraph): We split segmented objects that were assigned more than 25 synapses in the matching procedure into multiple pieces by applying a watershed operation with more conservative settings (foreground_threshold = 0.5, boundary_distance_threshold = 0.5, center_distance_threshold = 0.4, and no distance smoothing) to their masked area. This procedure was applied because large synapse counts are not physiological. We validated empirically that objects with these counts overwhelmingly corresponded to multiple IHCs wrongly merged into a single object. Note that this approach does not guarantee that all wrongly merged IHCs are separated. Even IHCs with a synapse count larger than 25 may not be split up if the watershed with more conservative setting still results in a single object.

For synapse detection, we trained the network to predict a single output channel that regressed a heatmap computed by convolving a binary map with an intensity of one at the annotated locations and zero otherwise with a Gaussian kernel with a sigma of one. We used the L2 loss to compare target and prediction. We used the same training hyperparameters (number of iterations, learning rate, etc.) as for the segmentation networks, with a validation split that consisted of 13.3% of the total training data. During inference, we applied a local maximum filter to the network predictions to detect ribbon synapses. This methodology was adapted from STACC (Jeremias & Pape, 2025), a recent method for spot detection. For application to whole cochleae, we mapped the synapse detections to segmented IHCs, by computing the distance of each detection to the closest object in the segmentation. We discarded detections with a distance of more than three µm to allow assigning synapses that were in areas not sufficiently stained in the Vglut3 channel, but to remove detections too far from an IHC to be a ribbon synapse.

The network architecture and training procedures were implemented in the torch_em Python library, which uses PyTorch internally. The segmentation code was implemented in an efficient way that scales to large volumes, making use of the elf Python library. Additional post-processing and analysis functionality was implemented with scikit-image (van der Walt *et al*, 2014) and SciPy (Virtanen *et al*, 2020). Segmentation results were stored in OME-Zarr and detection results in a csv table. They were uploaded to the S3 storage that also contained the image data and could then also be accessed in MoBIE for visual inspection.

### Segmentation and detection evaluation

We evaluated the SGN and IHC segmentations as well as the ribbon synapse detections against manual annotations; see also “Data annotation”. We computed the false positives (FPs), false negatives (FNs), and true positives (TPs). For the segmentations, these measures were computed by matching each annotation to the closest segmented object with a maximum distance of 1.9 micrometer. The number of matches corresponds to the TPs, the number of unmatched annotations to FNs, and the number of unmatched segmented objects to FNs. For the synapse detections, we followed a similar approach and matched points in predictions and annotations with a maximum distance of 1.1 micrometer. We then computed 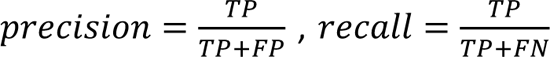, and 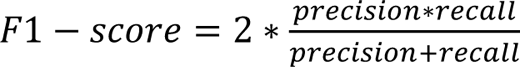 to measure the quality of results.

### Reference segmentation methods

We compared our segmentation methodology for SGNs and IHCs to popular general purpose cell segmentation methods: µSAM (Archit *et al*, 2025), using the light microscopy model, and CellPose (Stringer *et al*, 2021), using both CellPose v3 (Stringer & Pachitariu, 2025) and CellPose v4 (Pachitariu *et al*, 2025) models. In addition, we compared to Spiner (Cai *et al*, 2025), a dedicated method for the segmentation of SGNs in LSFM. Note that we did not retrain any of these models on our data, using the pretrained models provided by the respective tools. We also compared a StarDist (Schmidt *et al*, 2018) model trained on an earlier version of our SGN training data. Qualitative comparison of SGN and IHC segmentation results are shown for select methods in Figure 2. Quantitative evaluation and methodology are shown and explained in Ext. Data Figure 2.

We are aware of two other specific image analysis methods targeting the cochlea: HCAT (Buswinka *et al*, 2023b) and VASCilia (Kassim *et al*, 2025) for the segmentation and analysis of hair cells in confocal microscopy. However, HCAT only supports 2D segmentation and is thus not applicable to our data. While VASCilia can process 3D data, it mainly addresses stereocilia bundle analysis in hair cells and not hair cell segmentation.

### Quantification and analysis

We have implemented multiple quantification and analysis methods that address specific research questions based on segmentation and detection results.

For the tonotopic mapping of SGNs and IHCs (Figure 3a), we computed the run-length from base to apex by fitting a central path through the Rosenthal’s canal based on the 3D Euclidean distance transform of the center coordinates of the respective segmented cells. We then applied a Greenwood function according to (Ou *et al*, 2000) or (Müller *et al*, 2005) (Ext. Data Figure 5c) for mice and according to (Keppeler *et al*, 2021) for gerbil. It was applied to the relative run-length to obtain the frequency mapping. This function is parametrized as follows: f = A*10^(ax-k) with x being the fraction of the total distance between apex and base along the central path of the Rosenthal’s canal. The function of (Ou *et al*, 2000) corresponds to the parameter values A=1.46, a=1.77, and k=0, the mapping according to (Keppeler *et al*, 2021) corresponds to A=0.35, a=2.1, and, k=0.7, and the mapping according to (Müller *et al*, 2005) corresponds to A=1, a=1.21, and k=-0.68.

For the SGN (sub)type analysis (Figure 3, Ext. Data Figure 3), we first measured the median expression of the respective fluorescent (sub)type stains (Prph, CR, Ntng1, Calb1, Lypd1) as well as the median SGN intensity for each SGN soma. We then computed the ratio of median (sub)type stain intensity to median SGN stain intensity for each soma. Two annotators then determined the threshold at which this ratio best separated positive and negative expression of the respective stain for six small sub-volumes per cochlea (Ext. Data Figure 3c and d). We then determined positive / negative expression by applying the threshold of the spatially closest sub-volume. In some cases, where some sub-volumes in a given cochlea did not contain sufficient negatively or positively expressing cells, we instead determined a global ratio threshold. This approach was chosen to account for differences in staining perfusion. We then assigned a (sub)type to each SGN based on the stains it expressed, e.g. type I for cells not expressing Prph and type II for cells expressing Prph (see Results for details) to determine the overall fraction of (sub)types and their tonotopy.

For the evaluation of the ChReef therapy in mouse cochleae (Figure 4) and the evaluation of the f-Chrimson therapy in gerbil cochleae (Figure 5), we measured the median GFP expression per SGN after subtracting the GFP background signal in a radius of 75 µm. Two annotators then determined the threshold separating positive from negative expression for five (mouse) / seven (gerbil) sub-volumes per cochlea. We then determined the negative / positive expression for each SGN by applying the threshold of the spatially closest sub-volume. We measured the overall expression fraction as well as its tonotopy based on these values.

For the evaluation of the *OTOF* gene therapy in the samples from (Rankovic *et al*, 2021) (Figure 6c), we had to take a different approach for determining negative / positive expression in IHCs compared to the approach for SGNs described in the previous paragraph. This was due to a slight misalignment of the *rbOtof* channel (labeling transduction) and the CR channel (used for IHC segmentation), preventing direct computation of median intensity. Instead, we dilated the IHC segmentation by 2 pixels isotropically and then accumulated different statistics of the *rbOtof* intensities for each dilated IHC mask (mean, median, and standard deviation as well as the 1^st^, 10^th^, 25^th^, 75^th^, 90^th^, and 99^th^ percentile). We used these statistics as input features to a Random Forest Classifier, with labels for training derived from manual annotation of clearly positive / negative cells. This classifier was then applied to all IHCs to obtain the expression labels, which were used to compute the subsequent expression fractions and tonotopy.

## Data Availability

The data generated in this study will be made publicly available in a suitable data repository upon acceptance of the manuscript.

## Code Availability

The CochleaNet software is available at https://github.com/computational-cell-analytics/cochlea-net The version of the software at submission of this manuscript was 0.1.0. The software contains a Python library that implements the underlying functionality, a command line interface to apply segmentation and detection to large volumetric data, and a napari plugin to apply segmentation and detection to small volumes in a user-friendly manner. Its documentation is available at https://computational-cell-analytics.github.io/cochlea-net. The different neural networks provided by CochleaNet can be downloaded with the Python library. They will be deposited on BioImage.IO (Ouyang *et al*, 2022) upon acceptance of the manuscript.

## Extended Data Figures

**Extended Data Figure 1:**
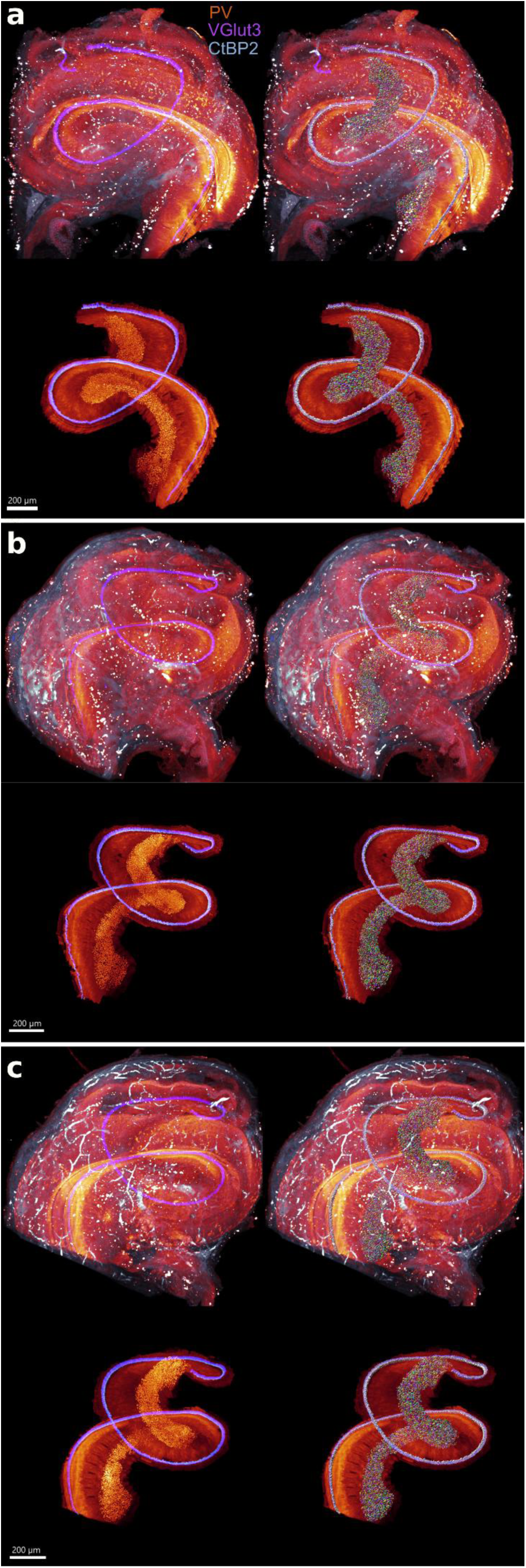
Three intact cochleae from untreated mice imaged with the high-resolution light-sheet microscope and analyzed with CochleaNet. a-c, Visualization of three cochlea with rendering of the unprocessed PV (red), Vglut3 (blue) and CtBP2 (white) signal (top left), the unprocessed signal with an overlay of SGN and IHC segmentation (top right), the processed signal, where off-target signal was removed and PV signal in SGNs as well as Vglut3 and CtBP2 signal in IHCs were increased to highlight the cells that were analyzed (bottom left), and overlay of the processed signal with segmentations (bottom right). Signal removal and intensity modulation was performed in Imaris using the segmentation masks from CochleaNet. Note that the CochleaNet networks were applied to the unprocessed signal; differentiation of signals in the cells of interest and background / off-target signal is one of the main analysis challenges.

**Extended Data Figure 2:**
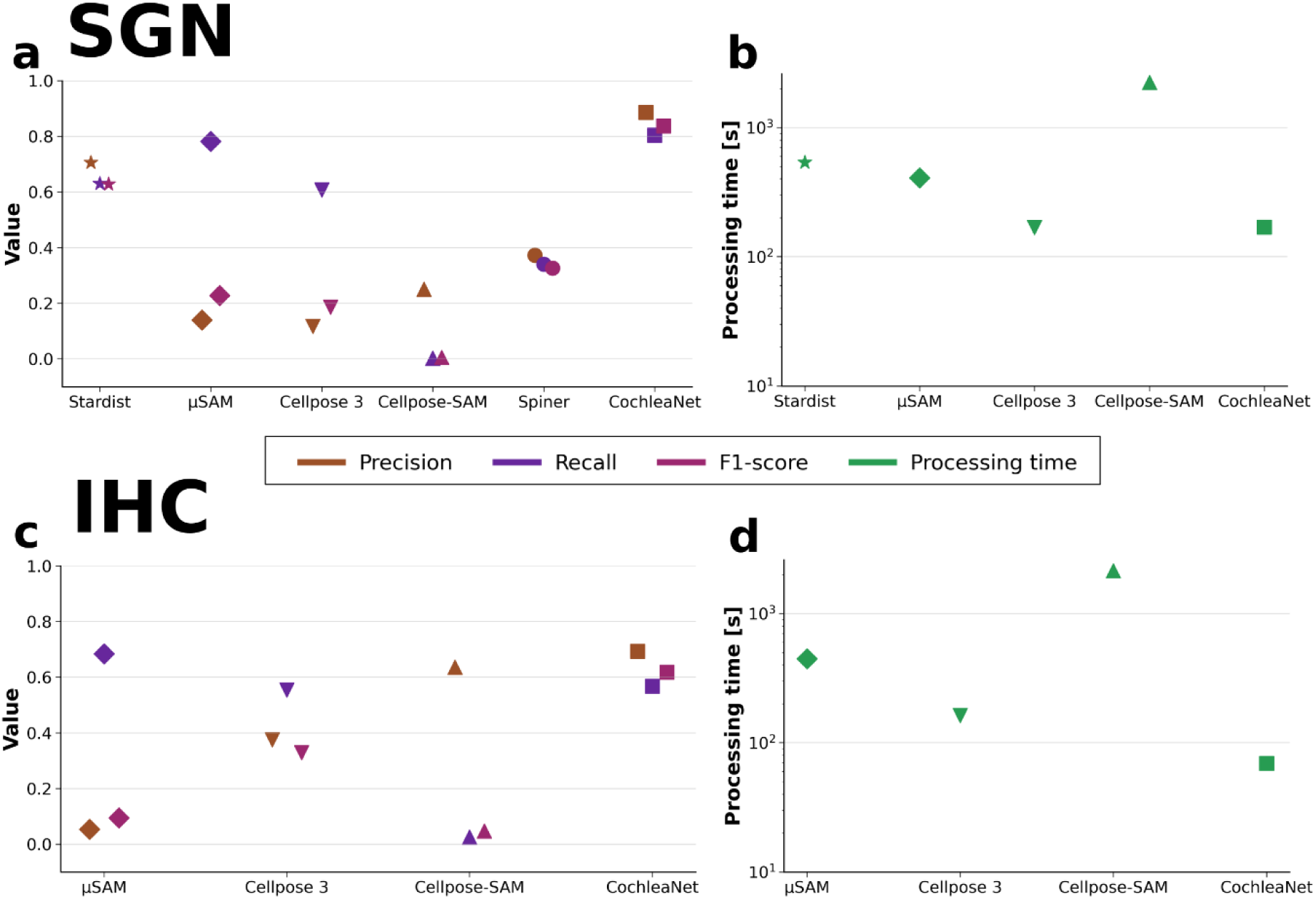
Extended quantitative comparison of segmentation methods. a, Evaluation of six different automated SGN segmentation methods against manual labels in twelve image planes. The methods were applied to a volume of ten planes above / below the central plane with annotations, except Spiner, which was applied only to the central plane as it is restricted to 2D segmentation. b, Average processing times (logarithmic scale) for segmenting those sub-volumes (a). We did not measure the processing time for Spiner, which was used in a GUI and applied only in 2D. c, Evaluation of four different automated IHC segmentation methods against manual labels in twelve image planes, following the same overall procedure as for SGNs. d, processing times corresponding to c. Overall, we see that CochleaNet yields clearly better results than pretrained generalist cell segmentation methods (Cellpose3, Cellpose-SAM, µSAM) and previously described specialized methods for the cochlea (Spiner), underscoring the need for a new, specialized solution. Moreover, it is more efficient than other methods, especially than Cellpose-SAM and µSAM, which is crucial for processing large volumetric data.

**Extended Data Figure 3:**
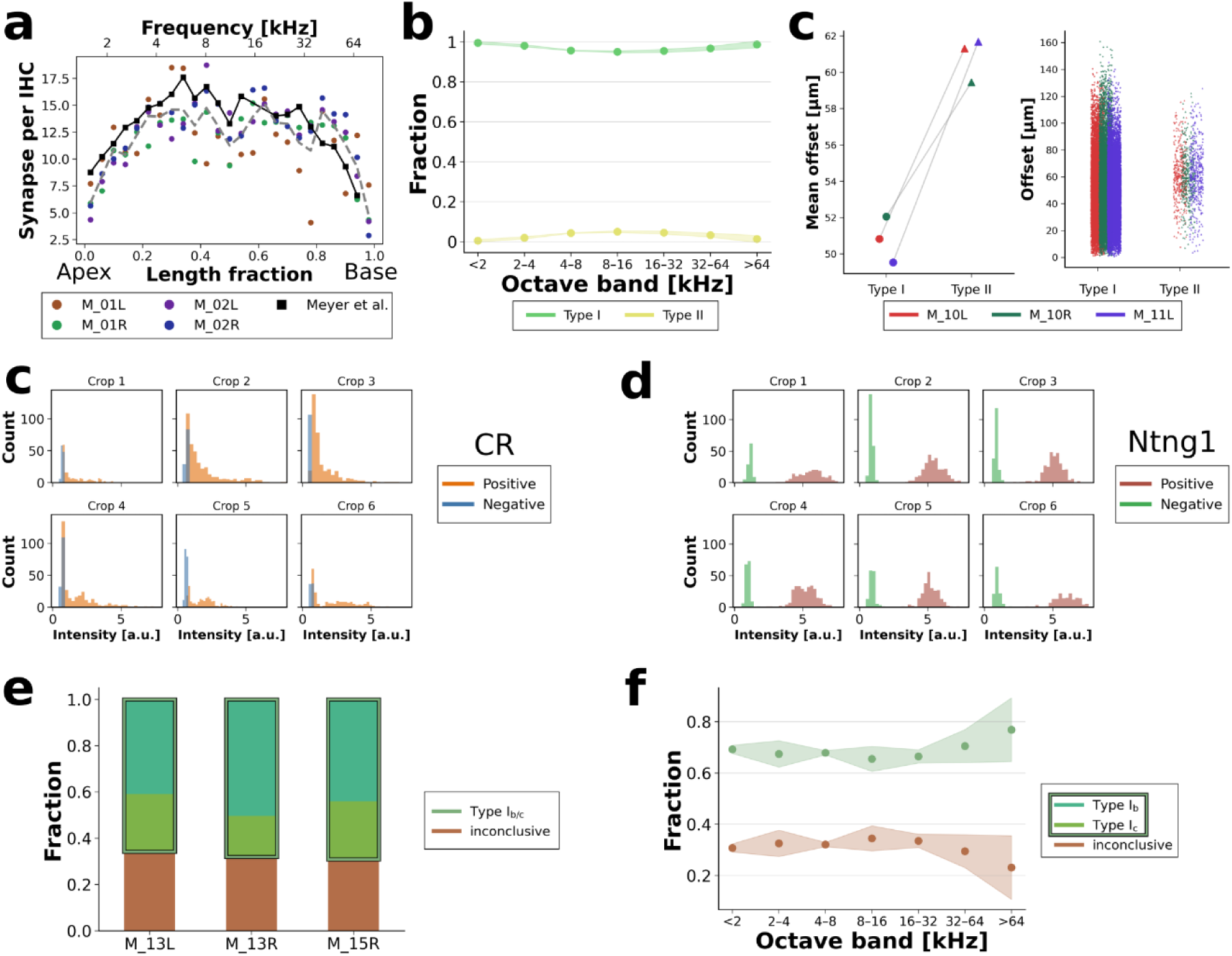
Extended analysis of synapse tonotopy and SGN (sub)types. a, Average synapse count per IHC of four intact cochleae (colored dots) compared to (Meyer *et al*, 2009) (black squares). Synapse counts are binned by 4% of the run-length, corresponding to approximately 200 micrometer of basilar membrane. Black line connects measurements from (Meyer *et al*, 2009), dotted grey line shows average of CochleaNet results. b, tonotopic mapping of SGN type I and type II for three cochleae with PV and Prph staining. c, Distance from central position of the ganglion for type I and type II SGNs (same cochleae as b). (Left) mean distance (right) scatter plot of individual soma distances. d, e, Histogram of the ratio of median CR-(c) and Ntng1-(d) to PV expression for the sub-volumes that were manually analyzed for determining expression thresholds. CR shows a continuous intensity distribution, corresponding to the different strengths of expression in different SGN subtypes, whereas Ntng1 shows a more binary expression pattern. f, fraction of subtype I_b_ and I_c_ SGNs determined in three cochleae with PV, Calb1, and Ntng1 staining. g, tonotopic mapping of subtype I_b_ and I_c_ for the data from f.

**Extended Data Figure 4:**
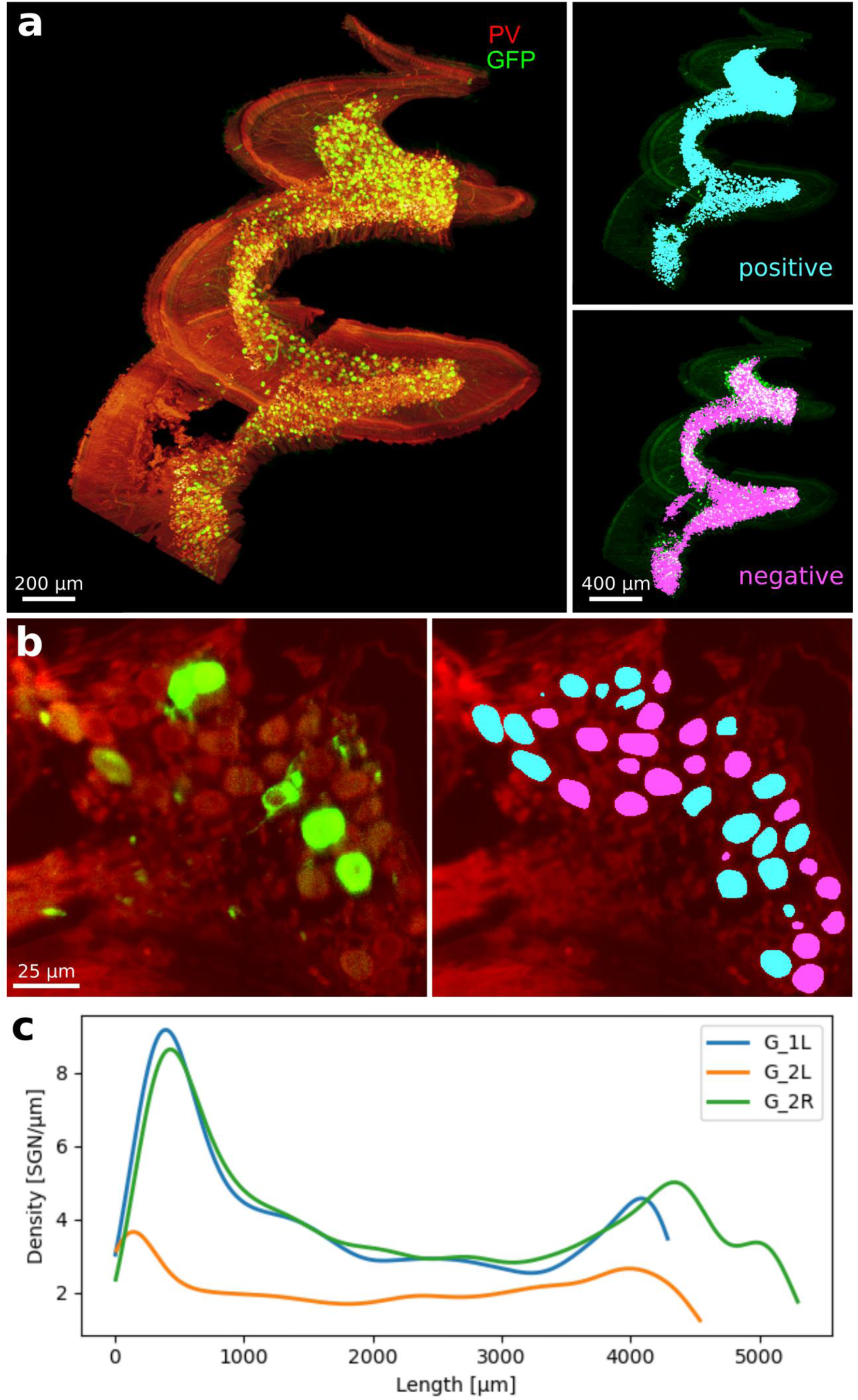
Extended analysis of gerbil cochleae. a, (Left) Rendering of the non-injected cochlea of an f-Chrimson treated gerbil showcasing gene therapy spread labelled with PV-(red) and GFP-(green) signal, the latter labelling f-Chrimson expression. (Right) overlay of SGN masks with positive (top, cyan) and negative (bottom, magenta) expression determined by our analysis. b, Zoom-ins of image signal (left) and corresponding expression masks (right). c, SGN density across the length of the Rosenthal’s canal for three cochleae, from an untreated gerbil (G1L) as well as left (injected, G2L) and right (non-injected, G2R) cochlea from an animal with postnatal optogenetic therapy for expressing f-Chrimson. Lower density in the injected cochlea (G_2L) indicates SGN loss due to the postnatal therapy.

**Extended Data Figure 5:**
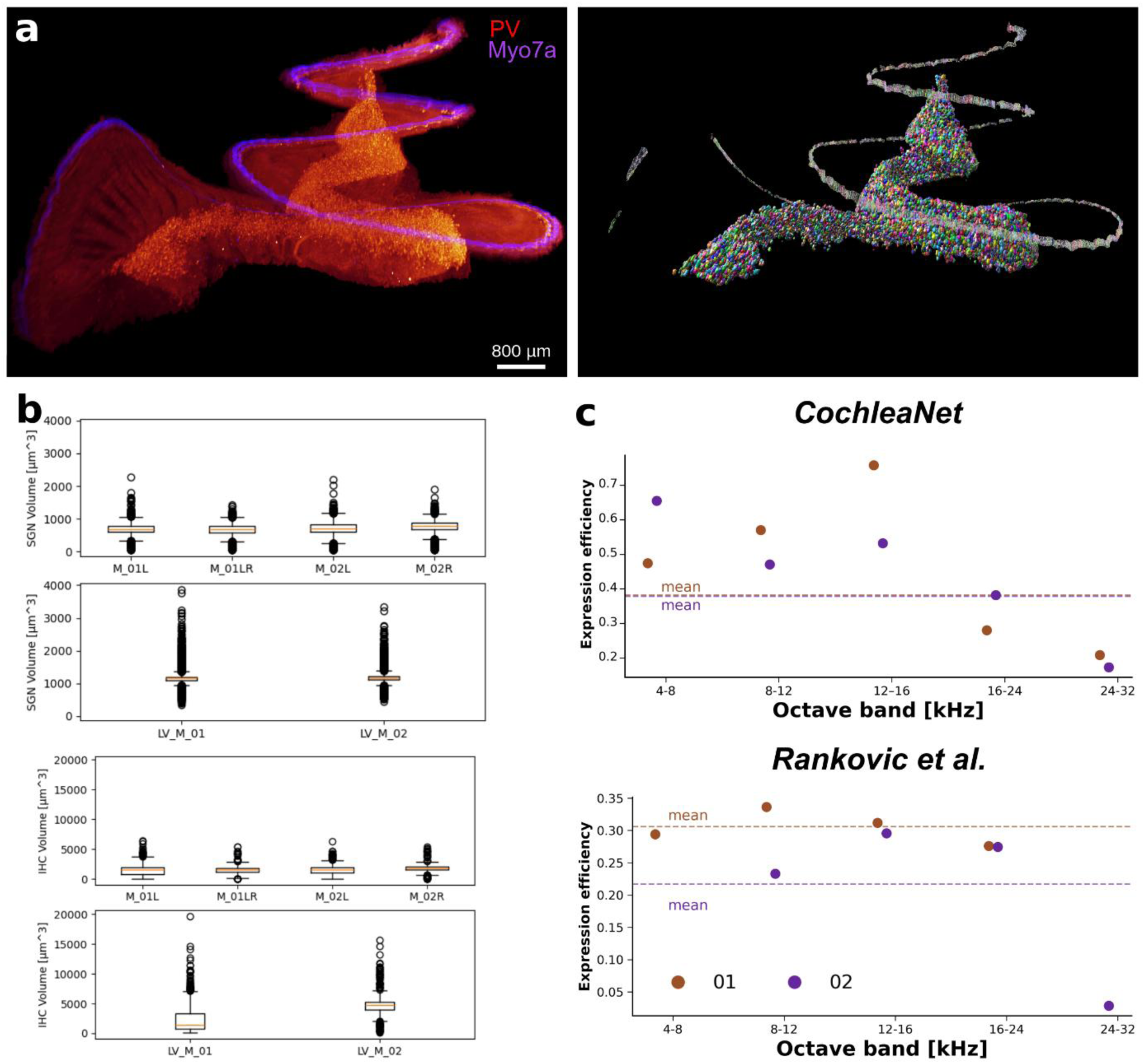
Extended analysis of cochleae imaged by a lower-resolution LSFM. a, Rendering of data of a marmoset cochlea taken from (Keppeler *et al*, 2021), showing PV (red) and Myo7a (blue) signal (left) as well as SGN and IHC segmentation (right). Note that the IHC segmentation contains gaps due to low signal-to-noise areas in the Myo7a signal. b, Comparison of murine SGN soma volumes derived from segmentations in four cochleae imaged with the high-resolution LSFM (top) and two cochleae imaged with the low-resolution LSFM (bottom). Higher average volume and larger number of outliers with high volume in measurements derived from low-resolution imaging indicate that cells were merged in the analysis of this data due to the difficulty of distinguishing closely packed cells, offering an explanation for the lower overall cell count compared to high-resolution imaging. Lower panels show the same analysis for IHCs, confirming the trend observed for SGNs c, Comparison of analyses of cochleae with *OTOF* gene replacement therapy imaged by low-resolution LSFM (Rankovic *et al*, 2021) and analyzed by CochleaNet (top) and manually (Rankovic *et al*, 2021) (bottom). CochleaNet estimated higher overall expression efficiencies (0.3820 and 0.3788 compared to 0.3059 and 0.2168). Note that CochleaNet analysis was based on more IHCs, 643 and 520 (we exclude IHCs from a torn-off component, hence the number is different compared to the one reported earlier) that were automatically segmented compared to 340 and 346 manually analyzed in (Rankovic *et al*, 2021). Tonotopic mapping according to (Meyer *et al*, 2009).

## Extended Data Tables

**Extended Data Table 1:**
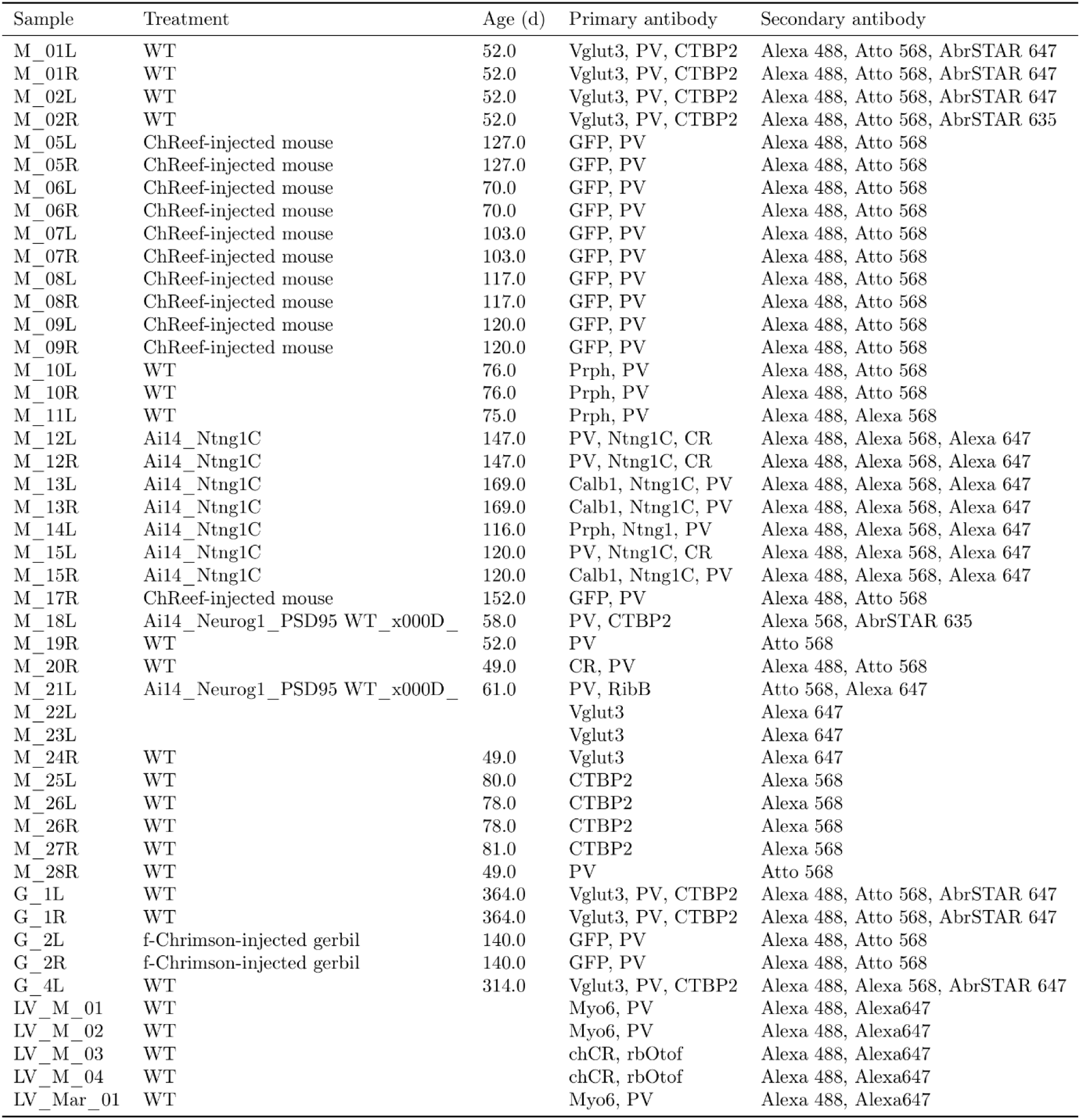
Overview of the cochleae used in this study, including the treatment applied, if any, the age, and the primary as well as secondary antibody used for fluorescent labelling; see Ext. Data Table 2 for details on labelling. Note that “LV” samples are re-used from previous studies (Keppeler *et al*, 2021; Rankovic *et al*, 2021); please refer to the respective publications for details.

| Sample | Treatment | Age (d) | Primary antibody | Secondary antibody |
| --- | --- | --- | --- | --- |
| M_01L | WT | 52.0 | Vglut3, PV, CTBP2 | Alexa 488, Atto 568, AbrSTAR 647 |
| M_01R | WT | 52.0 | Vglut3, PV, CTBP2 | Alexa 488, Atto 568, AbrSTAR 647 |
| M_02L | WT | 52.0 | Vglut3, PV, CTBP2 | Alexa 488, Atto 568, AbrSTAR 647 |
| M_02R | WT | 52.0 | Vglut3, PV, CTBP2 | Alexa 488, Atto 568, AbrSTAR 635 |
| M_05L | ChReef-injected mouse | 127.0 | GFP, PV | Alexa 488, Atto 568 |
| M_05R | ChReef-injected mouse | 127.0 | GFP, PV | Alexa 488, Atto 568 |
| M_06L | ChReef-injected mouse | 70.0 | GFP, PV | Alexa 488, Atto 568 |
| M_06R | ChReef-injected mouse | 70.0 | GFP, PV | Alexa 488, Atto 568 |
| M_07L | ChReef-injected mouse | 103.0 | GFP, PV | Alexa 488, Atto 568 |
| M_07R | ChReef-injected mouse | 103.0 | GFP, PV | Alexa 488, Atto 568 |
| M_08L | ChReef-injected mouse | 117.0 | GFP, PV | Alexa 488, Atto 568 |
| M_08R | ChReef-injected mouse | 117.0 | GFP, PV | Alexa 488, Atto 568 |
| M_09L | ChReef-injected mouse | 120.0 | GFP, PV | Alexa 488, Atto 568 |
| M_09R | ChReef-injected mouse | 120.0 | GFP, PV | Alexa 488, Atto 568 |
| M_10L | WT | 76.0 | Prph, PV | Alexa 488, Atto 568 |
| M_10R | WT | 76.0 | Prph, PV | Alexa 488, Atto 568 |
| M_11L | WT | 75.0 | Prph, PV | Alexa 488, Alexa 568 |
| M_12L | Ai14_Ntng1C | 147.0 | PV, Ntng1C, CR | Alexa 488, Alexa 568, Alexa 647 |
| M_12R | Ai14_Ntng1C | 147.0 | PV, Ntng1C, CR | Alexa 488, Alexa 568, Alexa 647 |
| M_13L | Ai14_Ntng1C | 169.0 | Calb1, Ntng1C, PV | Alexa 488, Alexa 568, Alexa 647 |
| M_13R | Ai14_Ntng1C | 169.0 | Calb1, Ntng1C, PV | Alexa 488, Alexa 568, Alexa 647 |
| M_14L | Ai14_Ntng1C | 116.0 | Prph, Ntng1, PV | Alexa 488, Alexa 568, Alexa 647 |
| M_15L | Ai14_Ntng1C | 120.0 | PV, Ntng1C, CR | Alexa 488, Alexa 568, Alexa 647 |
| M_15R | Ai14_Ntng1C | 120.0 | Calb1, Ntng1C, PV | Alexa 488, Alexa 568, Alexa 647 |
| M_17R | ChReef-injected mouse | 152.0 | GFP, PV | Alexa 488, Atto 568 |
| M_18L | Ai14_Neurog1_PSD95 WT_x000D_ | 58.0 | PV, CTBP2 | Alexa 568, AbrSTAR 635 |
| M_19R | WT | 52.0 | PV | Atto 568 |
| M_20R | WT | 49.0 | CR, PV | Alexa 488, Atto 568 |
| M_21L | Ai14_Neurog1_PSD95 WT_x000D_ | 61.0 | PV, RibB | Atto 568, Alexa 647 |
| M_22L |  |  | Vglut3 | Alexa 647 |
| M_23L |  |  | Vglut3 | Alexa 647 |
| M_24R | WT | 49.0 | Vglut3 | Alexa 647 |
| M_25L | WT | 80.0 | CTBP2 | Alexa 568 |
| M_26L | WT | 78.0 | CTBP2 | Alexa 568 |
| M_26R | WT | 78.0 | CTBP2 | Alexa 568 |
| M_27R | WT | 81.0 | CTBP2 | Alexa 568 |
| M_28R | WT | 49.0 | PV | Atto 568 |
| G_1L | WT | 364.0 | Vglut3, PV, CTBP2 | Alexa 488, Atto 568, AbrSTAR 647 |
| G_1R | WT | 364.0 | Vglut3, PV, CTBP2 | Alexa 488, Atto 568, AbrSTAR 647 |
| G_2L | f-Chrimson-injected gerbil | 140.0 | GFP, PV | Alexa 488, Atto 568 |
| G_2R | f-Chrimson-injected gerbil | 140.0 | GFP, PV | Alexa 488, Atto 568 |
| G_4L | WT | 314.0 | Vglut3, PV, CTBP2 | Alexa 488, Alexa 568, AbrSTAR 647 |
| LV_M_01 | WT |  | Myo6, PV | Alexa 488, Alexa647 |
| LV_M_02 | WT |  | Myo6, PV | Alexa 488, Alexa647 |
| LV_M_03 | WT |  | chCR, rbOtof | Alexa 488, Alexa647 |
| LV_M_04 | WT |  | chCR, rbOtof | Alexa 488, Alexa647 |
| LV_Mar_01 | WT |  | Myo6, PV | Alexa 488, Alexa647 |

**Extended Data Table 2:**
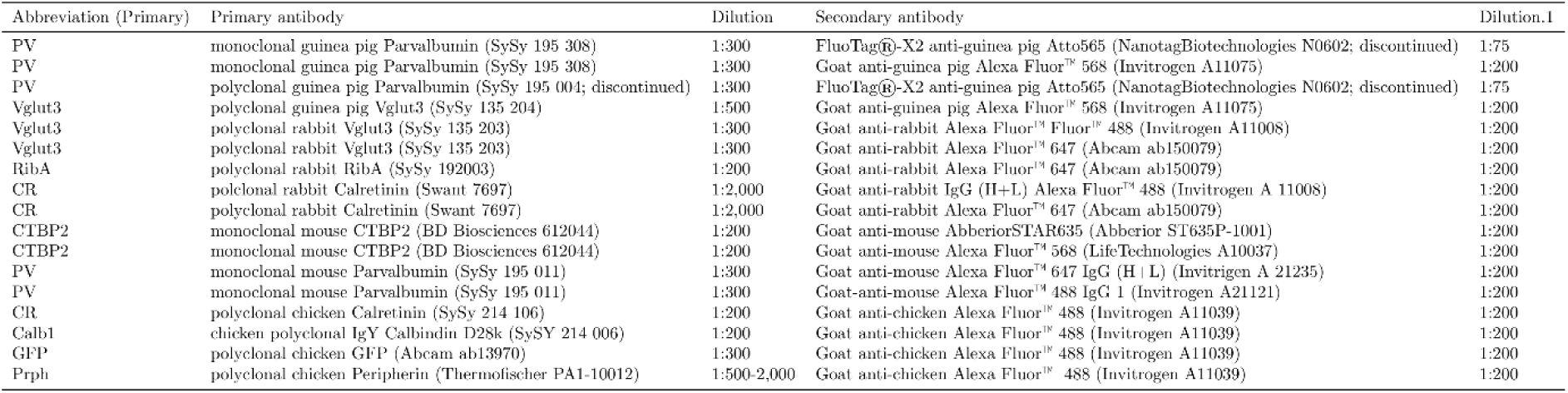
Overview of the different primary and secondary antibodies used in this study. This table lists all the respective combinations of primary and secondary antibodies as well as dilutions.

**Extended Data Table 3:**
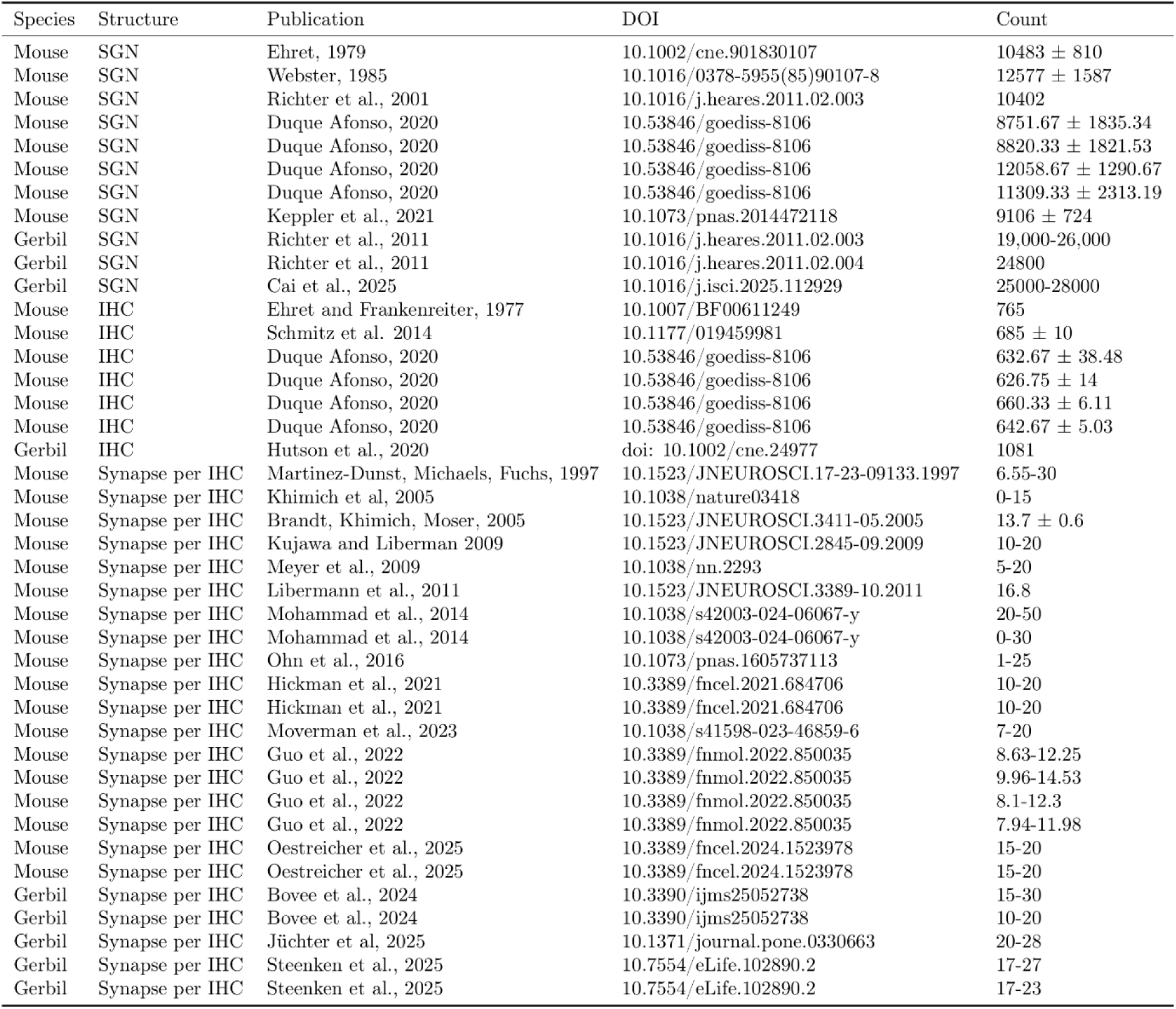
Summary of the literature values for SGN counts, IHC counts, and average count of ribbon synapses per IHC, in mouse and gerbil.

## Notes

https://github.com/computational-cell-analytics/cochlea-net

https://computational-cell-analytics.github.io/cochlea-net/

### Competing Interest Statement

T.M. is a co-founder of OptoGenTech company aimed at developing optogenetic hearing restoration.

